# An accurate and interpretable model for antimicrobial resistance in pathogenic *Escherichia coli* from livestock and companion animal species

**DOI:** 10.1101/2023.04.18.537400

**Authors:** Henri C. Chung, Christine L. Foxx, Jessica A. Hicks, Tod P. Stuber, Iddo Friedberg, Karin S. Dorman, Beth Harris

## Abstract

Understanding the microbial genomic contributors to antimicrobial resistance (AMR) is essential for early detection of emerging AMR infections, a pressing global health threat in human and veterinary medicine. Here we used whole genome sequencing and antibiotic susceptibility test data from 980 disease causing *Escherichia coli* isolated from companion and farm animals to model AMR genotypes and phenotypes for 24 antibiotics. We determined the strength of genotype-to-phenotype relationships for 197 AMR genes with elastic net logistic regression. Model predictors were designed to evaluate different potential modes of AMR genotype translation into resistance phenotypes. Our results show a model that considers the presence of individual AMR genes and total number of AMR genes present from a set of genes known to confer resistance was able to accurately predict isolate resistance on average (mean F_1_ score = 98.0%, SD = 2.3%, mean accuracy = 98.2%, SD = 2.7%). However, fitted models sometimes varied for antibiotics in the same class and for the same antibiotic across animal hosts, suggesting heterogeneity in the genetic determinants of AMR resistance. We conclude that an interpretable AMR prediction model can be used to accurately predict resistance phenotypes across multiple host species and reveal testable hypotheses about how the mechanism of resistance may vary across antibiotics within the same class and across animal hosts for the same antibiotic.

## Introduction

Antimicrobial resistance (AMR) is one of the foremost concerns in human and animal health and well-being. Multiple organizations including the World Health Organization, the European Commission, and major U.S. agencies such as the U.S. Centers for Disease Control and Prevention (CDC), the U.S. Food and Drug Administration (FDA), and the U.S. Department of Agriculture (USDA), have recognized the global threat of AMR infections [1–3]. AMR infections are estimated to cost more than US$20–35 billion per year in increased clinical treatment costs, and to contribute an additional 35,000 deaths per year in the U.S. alone [3, 4]. In response to this threat, the U.S. government has issued in 2015 the first National Action Plan for Combating Antibiotic-Resistant Bacteria (CARB NAP), directing all U.S. Federal agencies to collaboratively develop strategies against AMR infection using an integrated One Health approach [5]. The One Health approach recognizes the interconnection between humans, plants, animals, and the environment in global health, and emphasizes using information across sectors to understand and combat AMR [6]. CARB NAP includes the incorporation of multiple data streams such as the National Antimicrobial Resistance Monitoring System (NARMS), recommendation strategies for the judicious use of antibiotics in veterinary medicine, and restrictions on the use of certain glycopeptides, fluoroquinolones, and cephalosporins in both human and animal healthcare [7, 8].

As part of the USDA response to the CARB NAP, the Animal and Plant Health Inspection Service’s National Animal Health Laboratory Network (APHIS-NAHLN) established a collaborative AMR pilot project in 2018 with American Association of Veterinary Laboratory Diagnosticians (AAVLD) member laboratories [9]. A working group consisting of representatives from the AAVLD member laboratories, Clinical Laboratory Standards Institute (CLSI), the FDA Center for Veterinary Medicine Veterinary Laboratory Investigation and Response Network (VetLIRN), and the USDA

Center for Epidemiology and Animal Health (CEAH) was convened to recommend methods and standards that would meet the pilot project’s specific aims. These project aims included: (1) development of a sampling stream to monitor AMR in animal pathogens routinely isolated in veterinary diagnostic laboratories; (2) development of standardized methods for sharing and disseminating this information within the veterinary community, and (3) making data publicly accessible to inform future policies designed to mitigate AMR in animals [9]. Here we capitalized on the unique and extensive data from this pilot project to assess the state of knowledge and further elucidate the genetic mechanisms of AMR in this diverse set of agricultural and companion animals.

Genetic models of AMR are an important area of veterinary research. The decreased cost and increased availability of genome sequencing technologies has lead to a vast increase in the known genetic determinants of AMR, and the subsequent creation of over a dozen reference genetic databases and gene detection tools to assist AMR identification and research [10]. This information is then used to support diagnostic and surveillance observations made by traditional culture-based methods, and improve identification in cases where bacteria cannot be cultivated [11, 12]. Recently, researchers seeking to capitalize on this rapid growth of information have turned to machine learning (ML) models to further enhance AMR surveillance [13]. ML models have been successfully used to predict AMR resistance in pathogens such as non-typhoidal *Salmonella, Mycobacterium tuberculosis*, and *E. coli* [14–21]. These models use sequencing information to make accurate predictions of resistance phenotype, alleviating the need for antimicrobial susceptibility testing and potentially identifying new AMR genetic determinants in the process [22].

Due to this increasing interest, there is a large number of studies on genotype-to-phenotype relationships and ML-based prediction of AMR. However, there is little previous research on how these relationships vary depending on the specific antibiotic, animal host, or other factors. While several studies have found 95% to 100% correlation between AMR genotype and phenotype in multiple bacterial pathogens, these studies also find appreciable differences in the rates of resistance to fluoroquinolones, tetracyclines, aminoglycosides and *β*-lactams [23–25]. The differences in rates of resistance may be attributed to underlying differences in the resistance mechanisms of AMR gene products or may simply be due to some of them used exclusively in veterinary or animal husbandry practice [26, 27]. Additionally, bacterial isolates express widely variable rates of resistance depending on the host animal species [28, 29]. However, animal host information has rarely been considered in AMR models, and rarely do studies consider these relationships in more than one host animal at a time [30, 31]. Some differences in the micro-environments found in different animal hosts have been found to be a significant source of genetic variation [32–34]. This variation suggests that AMR genotype-phenotype relationships could be modeled in a more complex fashion. A meta-analysis examining the performance and reliability of ML models for AMR prediction in *Neisseria gonorrhoeae, Klebsiella pneumoniae* and *Acinetobacter baumannii* showed that model performance can vary significantly depending on the antibiotic, data set, resistance metric, and bacterial species [35]. Although ML methods in AMR prediction have been shown to be accurate, there is a need to better understand the variation in predictions and incorporate relevant biological and clinical knowledge into model design and evaluation before adoption in large-scale use.

Here we used sequenced isolates of *E. coli*, a zoonotic and cross-species pathogen, to indicate emerging AMR [36, 37]. *E. coli* may serve as a reservoir for transferring AMR genes to other animal and human populations through plasmids, transposons, and other mobile genetic elements [38]. Therefore, by examining the AMR profiles of *E. coli* isolates, we aim to develop an understanding of the different genetic associations that lead to the expression of AMR phenotypes in domesticated animals of interest to the clinical veterinary diagnostic community and public health practitioners.

We used genotype data from 980 *E. coli* isolates recovered from seven animal species to determine the utility of AMR genotype data for predicting phenotypic resistance in a clinically relevant context. Isolates came from various species of agricultural and companion animals as part of the APHIS-NAHLN AMR pilot project from 2018-2022. Our goal was to construct an interpretable, biologically informed model that would consider AMR gene content and host animal species to predict resistance to specific antibiotics. To do so, we fit and compared multiple elastic net models to characterize the relationship between AMR genes and predict resistance phenotype. We considered predictors that test different ways in which AMR genes could confer resistance. The proposed associations based on features that include the presence or absence of at least one gene within a group, the number of genes within each group, and the host animal effects. We then evaluated each model’s performance by its ability to predict AMR phenotypes. Our results show that while the different models have high predictive performance, there are significant differences in feature importance in predicting resistance to specific antibiotics in specific hosts. These differences suggest that future AMR modeling approaches could be made more accurate by incorporating information for each antibiotic and animal host. We discuss these differences in terms of the prevalence of resistance between different host animals, model complexity, gene association (binary presence/absence or summative count), and unique predictors. We also provide an estimate of our best fit model’s power to identify predictors of AMR phenotype in this study.

## Materials and Methods

The outline of this work is shown in Fig 1.

**Fig 1.**
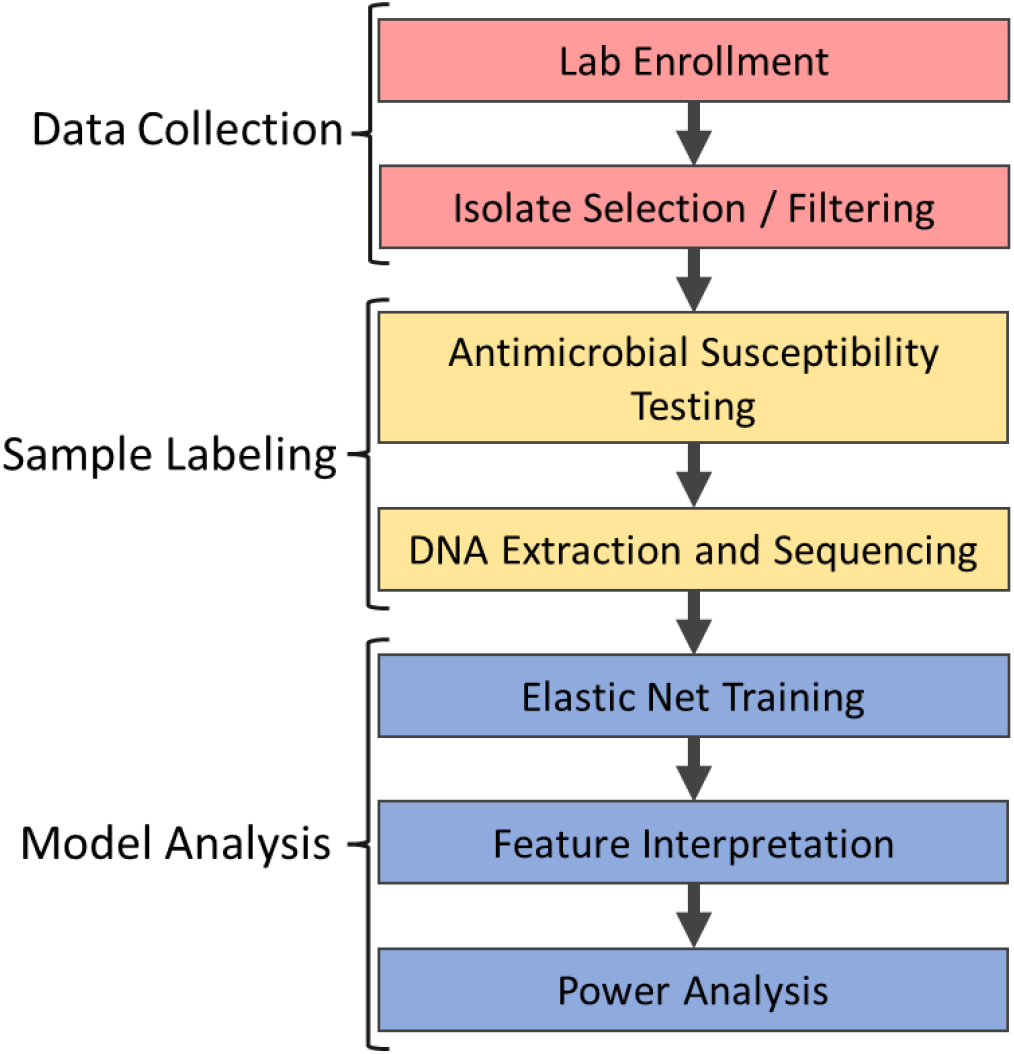
Project Workflow. We separate the project workflow into three distinct phases. In Data Collection (red); we enrolled participating USDA veterinary laboratories and collected *E. coli* isolates from animals associated with a clinical disease. We removed samples with improper or insufficient metadata. We then labeled samples (yellow) with their corresponding AMR phenotype and genotype using AST and WGS (see Methods). In the final model analysis (blue), we fit the phenotype and genotype information to multiple elastic net models, interpreted the importance of model features, and conducted a power analysis to assess the detectable effect sizes given the study sample sizes.

### Laboratory enrollment

Laboratories participating in the AMR pilot project were enrolled annually by APHIS-NAHLN. To maximize the representation of data at a national level, we considered factors such as animal population, representation, and geography in the enrollment process. Laboratories were enrolled and submitted the sequence data represented in this analysis. At the time of writing, the 27 participating laboratories were located in the following states: Alabama, California, Colorado, Florida, Georgia, Indiana, Iowa, Kansas, Kentucky, Louisiana, Michigan, Minnesota, Mississippi, Missouri, Nebraska, New York, North Carolina, North Dakota, Ohio, Pennsylvania, South Dakota, Texas, Washington, and Wisconsin.

### Isolate selection

We received data from 981 total *E. coli* samples from swine (102), cattle (198), chickens (121), turkeys (42), ducks (1), horses (151), dogs (195), and cats (173). Participating laboratories selected isolates for inclusion in the pilot study based on association with clinical disease, with one sample per distinct animal source (farm, herd, or owner within a single year). Each laboratory assigned a unique identifier to an isolate prior to submitting to APHIS-NAHLN, to eliminate all identifiable information from the data. Isolates were identified at the genus and species level by each laboratory, typically by using MALDI-TOF methods standard for veterinary microbiology laboratories [39].

Data submitted for each isolate included the minimum inhibitory concentration (MIC) values for all antibiotics tested, date of isolation, host animal species, specimen/tissue source, and final clinical diagnosis where available. After cleaning the data entries for typographical errors, we removed the duck sample from subsequent analyses as it represented a single animal from that species. Tissue samples were originally collected for other clinical purposes.

### Antimicrobial susceptibility testing

Participating laboratories conducted all *E. coli* antimicrobial susceptibility testing (AST) using the following commercially available Sensititre™plates (Thermofisher Scientific, USA) according to the manufacturer’s instructions: BOPO6F or BOPO7F (swine and cattle), EQUIN1F or EQUIN2F (horses), AVIAN1F (chickens and turkeys), and COMPGN1F (dogs and cats). The six plate layouts previously listed include a total of 50 antibiotics from 14 different antibiotic classes, with each plate type containing between 18-24 different antibiotics.

We used current clinical breakpoint guidelines from the Clinical and Laboratory Standards Institute (CLSI) established based on a combination of factors including the host animal species, sample source and type (*e*.*g*., urinary tract or skin/soft tissue infection), and bacterial isolate. Seventeen antibiotics have interpretive clinical breakpoints established for *E. coli* in various animal species at the time of writing in the *Vet01S* [40]. These were: amikacin (dogs, horses), amoxicillin-clavulanic acid (cats, dogs), ampicillin (cattle, cats, dogs, horses), cefazolin (dogs, horses), cefovecin (cats, dogs), cefpodoxime (dogs), ceftazidime (dogs), ceftiofur (cattle, swine), cephalexin (dogs), doxycycline (horses), enrofloxacin (cats, dogs, horses, poultry), gentamicin (dogs, horses), marbofloxacin (cats, dogs), minocycline (horses), orbifloxacin (cats, dogs), piperacillin-tazobactam (dogs), and pradofloxacin (cats, dogs). Based on these breakpoints, we classified samples as expressing sensitive, intermediate, or resistant antimicrobial phenotypes (Table S1). To simplify model prediction, intermediate phenotypes were labeled as antibiotic resistant.

For antimicrobials without CLSI breakpoints, samples were classified into wildtype (*wt*) and non-*wt* groups using epidemiological cut-off values (ECOFF) from the European Committee on Antimicrobial Susceptibility (EUCAST) [41]. Isolates with MIC values at or below the ECOFF value are considered to be members of the wildtype population for that given bacterial species and antimicrobial. ECOFF values are calculated by fitting a cumulative log-normal distribution using non-linear least squares regression to MIC data curated by the EUCAST database [42]. ECOFF values were collected from the EUCAST MIC distribution website (http://www.eucast.org) in February 2023. Unlike CLSI breakpoints, ECOFF values are non-host animal specific. If an isolate’s phenotype was ambiguous, such as MIC values reported as “less than or equal” to a value exceeding its CLSI breakpoint or ECOFF value, the sample phenotype was labeled “Non-Interpretable”. Isolate and antimicrobial combinations without a CLSI breakpoint or ECOFF value, or with a Non-Interpretable phenotype, were removed from further analysis (Table S2).

### DNA extraction and whole-genome sequencing

Isolates were either sequenced directly by participating laboratories or submitted to the National Veterinary Services Laboratories (NVSL) for whole-genome sequencing (WGS). Briefly, extracted DNA was used to prepare indexed genomic libraries using the Nextera XT^®^ DNA Library Prep Kit (Illumina). Multiplexed libraries were sequenced using 250 × 2 paired-end read chemistry on the Illumina MiSeq^®^ platform at an average sequencing depth of 91.2 ± 32.3-fold.

Isolates were verified as *E. coli* using Kraken against a database consisting of RefSeq complete genomes database release ver. 2.09 [43], UniVec-core, and host genomes commonly encountered by the NVSL, with 85% reference genome coverage or greater after Bayesian Reestimation of Abundance, using default parameters [44, 45]. Following this validation step, *de novo* assembly of bacterial genomes was performed using SPAdes v3.14.0 [46] and resultant scaffolds were processed using the AMRFinder v1.0.1 [47] and ABRicate v3.2.3 [48] tool kits against the National Center for Biotechnology Information (NCBI), AMRFinder, and ResFinder [49] databases. Resultant gene hits from these tool kits, with minimum inclusion requirements of 70% amino-acid identity and 95% reference gene coverage, were then collated with the metadata as putative AMR genes found in each *E. coli* isolate. Plasmid genes were not used due to insufficient information linking the presence or absence of those genes with specific antibiotic resistance. In total, 197 different AMR genes were identified across all samples.

### Determining animal host differences

We used Fisher’s exact test to determine if there were significant differences in the fraction of resistant or non-*wt* phenotypes between animal hosts for each antibiotic in the data set. We applied the Bonferroni-Holm (BH) correction for multiple comparisons to *p*-values, and a BH corrected *p*-value threshold of 0.05 was used to determine significance. All analyses and statistical tests were done using R 4.2.2, model training and evaluation was performed using the tidymodels family of packages, and visualizations created with the ggplot2 3.4.1 [50–52].

### Elastic net regularization

We used an elastic net model to estimate the effect of relevant genotype predictors on the resistance phenotype. An elastic net is a penalized regression model which combines the penalties of Least Absolute Shrinkage and Selection Operator (LASSO) and Ridge regressions [53–55]. The elastic net utilizes two tunable parameters, a shrinkage parameter *λ* for the LASSO regression, and a mixing parameter *α* that determines the relative importance of the LASSO and Ridge penalties when they are added together. Rifge-driven shrinkage and LASSO-driven predictor selection make elastic net suitable for AMR prediction from genotype, where the number of predictors greatly exceeds the number of samples, while retaining a higher level of predictor interpretability compared with other ML methods [56]. The candidate elastic net predictors were built from relevant antibiotic genes, defined as the set of AMR genes known to cause resistance to the specified antibiotic.

We curated a custom database that captures the knowledge of gene-specific AMR based on common published literature (See Data Availability). In this database, AMR genes are associated with specific individual antibiotics or a general class of antibiotics. Throughout this analysis, we separately fit samples with CLSI and ECOFF breakpoints. In addition, to avoid fitting models with low power, we excluded antibiotics represented in fewer than 5 samples per phenotype label. For the remaining 31 antibiotic and breakpoint combinations, we only admitted genes into the relevant set χ_*i*_ that are known to confer resistance to antibiotic *i*. The exception was the *Full* model, where all AMR genes, including genes associated with other antibiotics, were considered. We then used elastic net with the logistic regression (Equation (1)) to model and predict isolate antibiotic resistance based on the presence or absence of genes in the gene group.

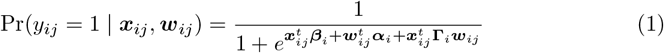

In equation (1), *y*_*ij*_ is an element in the binary vector 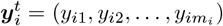, where *y*_*ij*_ = 1 denotes resistance or non-*wt* response to the *i*-th of *n* antibiotics in the *j*-th of *m*_*i*_ samples. Vector 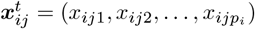, where *x*_*ijk*_ *∈ {*0, 1*}* represents the presence or absence in the *j*-th sample of the *k*-th of *p*_*i*_ predictors which confer resistance or non-*wt* phenotype to the *i*th antibiotic. Vector 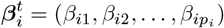 denotes the set of regression coefficients for the gene effects to antibiotic *i*, each *β*_*ik*_ a measure of the relative degree of association between a gene *k* and the resistance or non-*wt* phenotype to antibiotic *i*. To account for potential effects of the animal host on the rate of isolate response, we include a set of predictors, 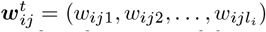 corresponding to a one-hot encoding of the animal host for each isolate *j* tested for resistance or non-*wt* phenotype to antibiotic *i* and coefficients 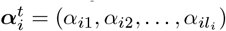 for the effect of each animal on resistance or non-*wt* phenotype to antibiotic *i* among *l*_*i*_ distinct species treated with this antibiotic. For example, in a model for antibiotic *i* with data only from swine and cattle samples, ***w***_*ij*_ is the length-two predictor for “swine” and “cattle” with 0 or 1 values to indicate if sample *j* came from a swine or cattle host. An interaction term, 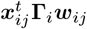, represents the animal-specific differences in the genetic effects for resistant or non-*wt* phenotypes in different animal hosts, with **Γ**_*i*_ a *p*_*i*_ ×*l*_*i*_ matrix of coefficients. To keep notation simple, we neglect adding an extra index for CLSI or ECOFF breakpoint, but some antibiotics participate in two fits with distinct estimates for each type of breakpoint. We also considered the possible effect location, the US state or region from which a sample was sourced, may have on phenotype. We found location and host animal labels were not independent (*χ*^2^ *<* 2.2 ×10^*−*16^), likely due to a bias for certain animal hosts to be raised in specific geographic regions, and we therefore only considered animal hosts as a relevant predictor.

To consider the relative predictive power of different predictors, we built models from sets of predictors implying distinct biological mechanisms of resistance, (Table 1), by default allowing main and gene interaction effects for host species. *Main effects* are those contributed by individual predictors while *interaction effects* are those in which the effect of a predictor is dependent upon one or more other predictors. The *Gene* model considers presence/absence of any gene in the relevant gene set χ_*i*_ known to confer resistance to the *i*-th antibiotic. The *Binary* model considers a single binary indicator *v*_*ij*_ = 1 if 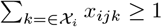, which will be predictive if the presence of any one gene from the relevant gene set χ_*i*_ is capable of conferring resistance. The *Count* model includes only the number 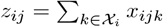 of observed relevant resistance genes as a predictor, implying each additional relevant gene from χ_*i*_ linearly increases the log odds of resistance. We also fit various combinations of these models. Lastly, we tested a *Full* model using all AMR genes identified in an isolate, ignoring biological knowledge about whether those genes are known to confer resistance to the predicted antibiotic. We define candidate predictors as the set of predictors used in each model.

**Table 1.** Elastic net model formulations. We use figurative equations to represent the different models, using *i* to index the current antibiotic in question. Covariates ***w***_*i*_ indicate host animal, ***x***_*i*_ indicate presence/absence of genes in the relevant set *χ*_*i*_ (***x*** for all AMR genes, not just those in the relevant set), ***v***_*i*_ indicate presence/absence of *any* relevant gene, and ***z***_*i*_ the count of observed relevant genes. Operator indicates interactions considered, while + indicates main effects only. See text for a detailed explanation of each model.

| Model | Formulation | Brief explanation |
| --- | --- | --- |
| Gene | $y_i \sim w_i * x_i$ | Is the relevant set ($\mathcal{X}_i$) of AMR genes predictive of resistance? |
| Gene & Binary | $y_i \sim w_i * (x_i + v_i)$ | Does knowing any gene in a given set in addition to the relevant gene set ($\mathcal{X}_i$) improve resistance prediction? |
| Gene & Count | $y_i \sim w_i * (x_i + z_i)$ | Does knowing the total number of genes (rather than individual identity) in addition to the relevant gene set ($\mathcal{X}_i$) improve resistance prediction? |
| Gene & Binary & Count | $y_i \sim w_i * (x_i + v_i + z_i)$ | A combination of the two previous models. |
| Full | $y_i \sim w_i * x$ | Is the total detectable set of AMR genes, ignoring known conferred resistance, a better predictor of resistance? |
| Count | $y_i \sim w_i * z_i$ | Is the total number of genes in the relevant set ($\mathcal{X}_i$) predictive of resistance? |
| Binary | $y_i \sim w_i * v_i$ | Is the presence of any gene within the relevant set ($\mathcal{X}_i$) predictive of resistance? |

### Model training and feature stability

After applying the inclusion criteria, we were left with 31 subsets of data meeting our requirements: 15 using CLSI breakpoints and 16 using ECOFF values, representing 24 unique antibiotics in total. We then fit separate elastic net models to each subset of data. For each model, we further split the subset of data into 80:20 training and testing splits, and fit the model with the training data. Within the training subset, a 10-fold cross validation scheme was used to select the value of the *α* and *λ* hyperparameters, optimizing the accuracy of predicted resistance phenotype. We evaluated the model fit by calculating the prediction accuracy of the fitted model with the testing data. When the number of candidate predictors is large, selected predictors for models can be unstable and vary between model fits [57, 58]. To identify important predictors, we used a stability selection method where we repeatedly refit the model (80:20 cross-validation and selection of *α* and *λ*) with a random 80% of candidate predictors over 1000 replicates [59, 60]. We considered important predictors to be selected predictors found in over 66% of model replicates. After training and stability selection, 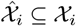 denotes the set of important predictors consistently included in the model.

### Model performance and power analysis

To evaluate model performance, we used a suite of evaluation metrics standard to binary classification models described in Table 2. Briefly, these different metrics discriminate the performance of the model to accurately identify samples sensitive or resistant to antibiotics, while considering the samples that are incorrectly predicted to be sensitive or resistant.

**Table 2.** Model Performance Metrics. Model performance was evaluated using F_1_ score, Negative Predictive Value (NPV), Positive Predictive Value (PPV), Sensitivity/Recall, and Specificity. The F_1_ score represents the harmonic mean of the Precision and Recall; NPV is a calculation of true negatives / all negatives; PPV is a calculation of true positives / all positives; Sensitivity/Recall is a calculation of true positives / (all true positives and false negatives); and Specificity is a calculation of true negatives / (all true negatives and false positives). TN, FN, TP, and FP refer to true negatives, false negatives, true positives, and false positives respectfully.

| Metric | Model |
| --- | --- |
| Negative Predictive Value (NPV) | $\frac{TN}{TN+FN}$ |
| Positive Predictive Value (PPV or Precision) | $\frac{TP}{TP+FP}$ |
| Sensitivity (Recall) | $\frac{TP}{TP+FN}$ |
| Specificity | $\frac{TN}{TN+FP}$ |
| F<sub>1</sub> Score | $2 \times \frac{\text{precision} \times \text{recall}}{\text{precision} + \text{recall}}$ |

There is no accepted method of power calculation for elastic net models. Instead, we sought to assess the *post hoc* power to detect a pre-specified effect size for each predictor in the *Gene&Binary&Count* model. Our goal was to, at least roughly, assess whether the observational study could have detected a moderate effect of each predictor given the observed sample sizes and assuming no interaction with other predictors. Whether a predictor actually gets selected into the fitted elastic net model, however, depends on the complicated relationships between predictors [61], which was ignored in our power analysis. We based our analysis on simple logistic regression and a Wald test-derived procedure described in [61] and implemented in the R *WebPower* package [61, 62] with the `wp.logistic()` function. We set *α* = 0.05 and distribution family to “Bernoulli” when testing binary predictors and to “normal” when testing the “count” predictor in relevant models. In all cases, we sought the power of detecting effect sizes of *β*_1_ = 0.7, 1.1, 1.6, 2.3 or a 2, 3, 5 or 10-fold increase in the odds of resistance upon inclusion (or unit increase) of the predictor. For the binary predictor *x*_*ijk*_ indicating the presence of gene *k* conferring resistance to antibiotic *i*, we set the null model resistance probability (parameter p0) to the observed proportion in the subset of observations *j* with *x*_*ijk*_ = 0 and the alternative model resistance probability (parameter p1) was set to the expected proportion of resistance upon the desired fold increase in odds of resistance. For the “count” predictor, we set p0 to the observed proportion of resistance in the sample and p1 to the proportion obtained upon increasing the odds of resistance by the desired fold.

## Results

We examined the prevalence of antibiotic resistant or non-*wt* phenotypes across 980 *E. coli* isolates. When evaluating prevalence, we only consider phenotypes with at least five samples. The highest prevalence of resistance was for minocycline (100%; *n* = 21), doxycycline (94.5%; *n* = 52), and ampicillin (92.5%; *n* = 571). The lowest prevalence was for amikacin (6.45%; *n* = 22) and piperacillin-tazobactam (6.74%; *n* = 13). Non-*wt* phenotypes were most prevalent against imipenem (66.7%; *n* = 4), ampicillin (52.6%; *n* = 91) and amoxicillin (35%; *n* = 57). We observed zero isolates with non-*wt* phenotypes against amikacin (*n* = 175). The following antibiotics had non-interpretable MIC values; azithryomycin (*n* = 116), cephalexin (*n* = 152), and florfenicol (*n* = 65).

Previous studies have shown isolate resistance varies between different animal species [28, 29, 63–65], and we confirm isolates from distinct animal hosts exhibit widely varying rates of AMR. We used a Fisher’s exact test to identify significant differences in the proportion of resistant isolates between host species for multiple antibiotics, with respect to CLSI or ECOFF values (Fig 2). For streptomycin and neomycin, the rate of resistance was significantly lower in chicken relative to isolates from other host animals. For gentamicin, another aminoglycoside, dog isolates had the lowest rate of resistance. Horse isolates had the largest differences in resistance against three drugs relative to other animal hosts, with over 85% of isolates resistant to cefazolin, ampicillin, and enrofloxacin. Cat, horse, and cattle isolates shared high rates of resistance to ampicillin, significantly greater than dog isolates. Although ECOFF values are not animal specific, we observe significant differences in the fraction of non-*wt* phenotypes for trimethoprim-sulphamethoxazole, chloramphenicol, tetracycline and doxycycline. Across shared antibiotics, turkey isolates had an overall higher proportion of resistant phenotypes than chicken isolates, and isolates from dogs had higher rates of resistance than isolates from cats. Even within similar groups of animals (e.g. avian species: chicken and turkey, or companion species: cats and dogs), there are significant differences in rates of resistance observed for a given antibiotic.

**Fig 2.**
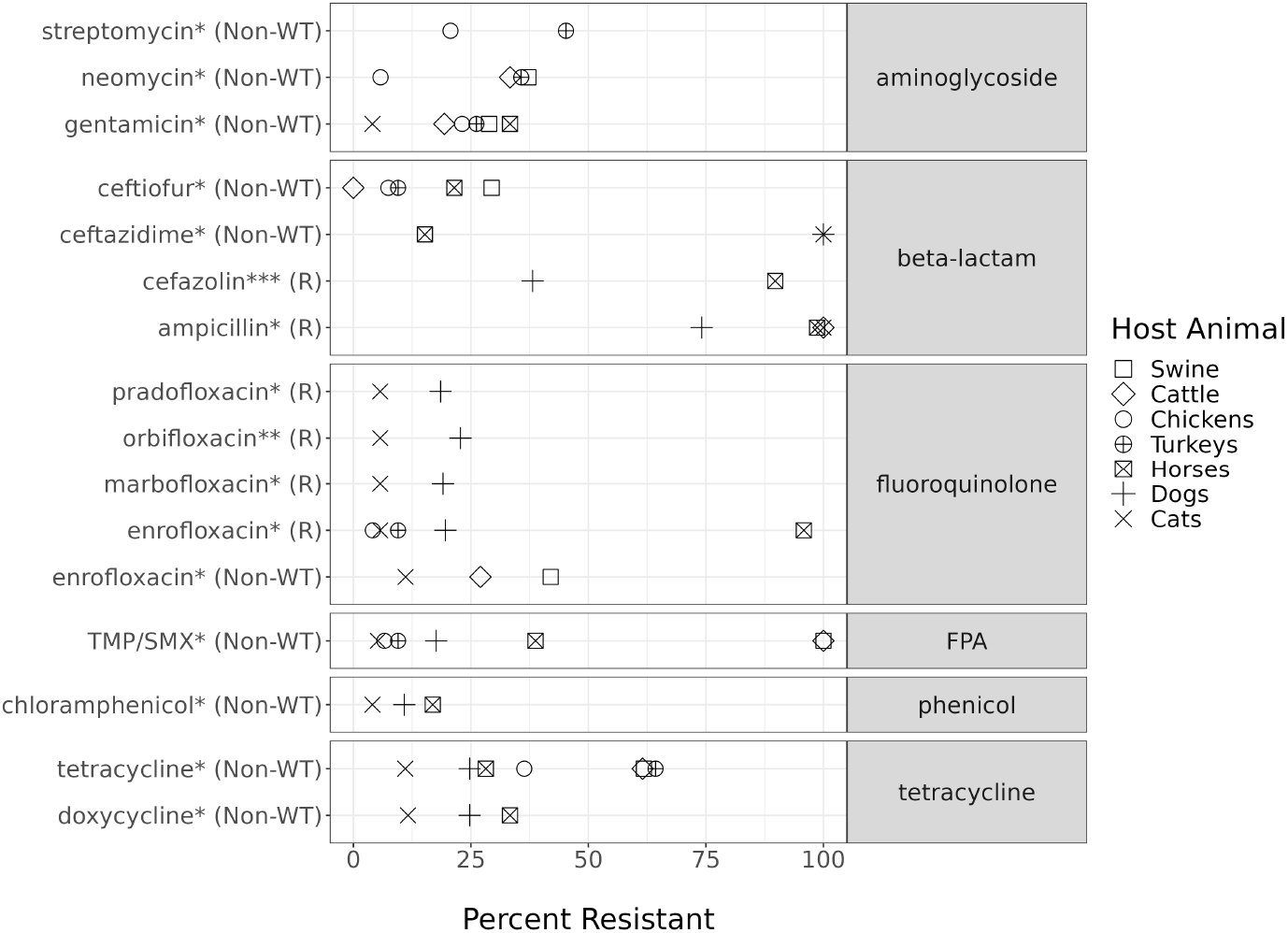
Significant differences in antibiotic resistance between animal hosts. The *y*-axis labels indicate the antibiotic and breakpoints used to determine resistant phenotype. The *x*-axis represents the percentage of resistant isolates. Symbols indicate animal host. Each point is the proportion of resistant isolates to the named antibiotic among those extracted from the indicated animal host. Only antibiotics with a significant difference in the proportion of resistance isolates between animal species are shown (adjusted *p*-value, *\* ≤* 0.05, *\*\* ≤*0.01, *\* * * ≤* 0.001). FPA = Folate Pathway Antagonist, TMP/SMX = trimethoprim-sulphamethoxazole.

### Predicting resistance

We fit increasingly complex elastic net models corresponding to types of association for the genetic determinants of AMR (Fig 3). We assume genotype-to-phenotype associations will vary depending on the host, so each model includes host animal main and interaction effects as a baseline. Many existing AMR models predict resistance based on the presence of *any* AMR genes relevant to the modeled antibiotic [66–68]. A relevant AMR gene is a gene known to confer resistance to a specific (or response) antibiotic. We fit an equivalent model, *Binary*, and measure its F_1_ score, or the harmonic mean of the precision and recall at 69.2% ±24.1%. We subsequently fit more complex models building upon reasonably assumed mechanisms of resistance. The *Count* model, which evaluates whether the log odds of resistance linearly increases with the presence of every additional AMR gene, achieved a mean F_1_ of 92.0% ±10.3% across all antibiotics. The *Gene* model uses the presence of individual AMR genes as separate predictors, allowing different genes to have distinct roles in conferring resistance. The *Gene* model, despite having more predictors, only achieved a F_1_ of 84.1% ±18%, lower than the *Count* model.

**Fig 3.**
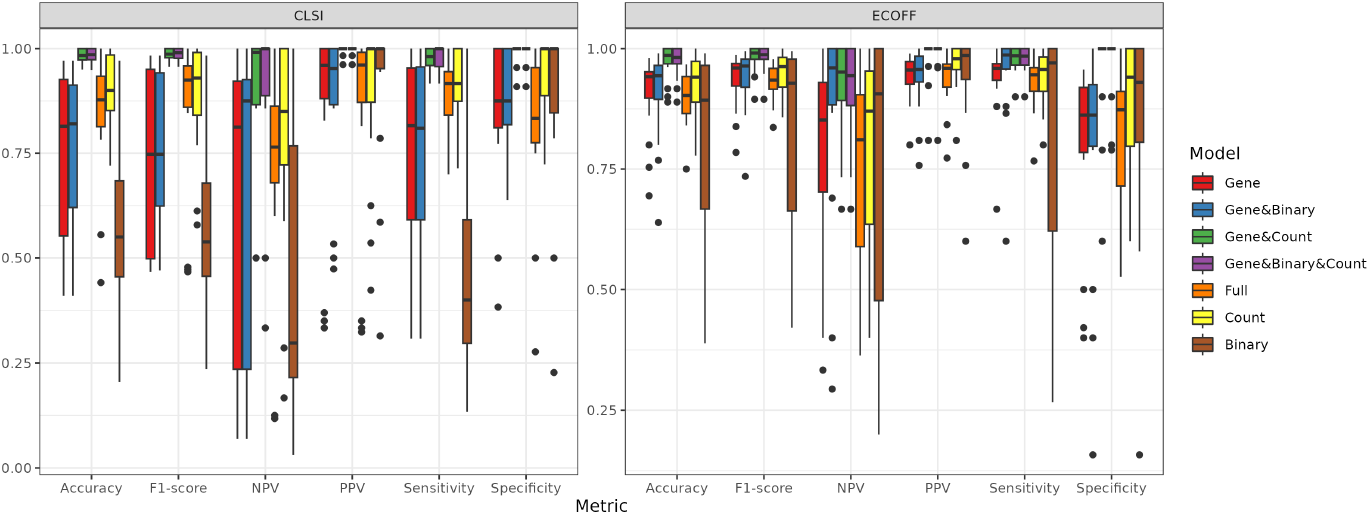
Elastic net model evaluation. We fit seven different elastic net models evaluating the effect of different predictors described in Table 1, including: AMR genes, animal host, presence of *n* ≥ 1 AMR gene (“binary”), and total counts of relevant AMR genes (*n*, “count”). The *Gene&Binary&Count* model best predicts the resistance phenotype, with the highest median F_1_ score and median accuracy across the different antibiotics. Model performance was evaluated using F_1_ score, NPV, PPV, Sensitivity/Recall, and Specificity. NPV = Negative Predictive Value; PPV = Positive Predictive Value.

We separately added the “binary” or “count” predictors to the *Gene* model, allowing the underpowered data sets to use simpler predictors in addition to individual genes. The *Gene&Binary* model achieved a slightly higher mean F_1_ of 85.5% *±* 16.1%. The *Gene&Count* achieved the best performance with a mean F_1_ of 98.2% *±* 2.3%, with a similar performance to that of the *Gene&Binary&Count* model with a mean F_1_ of 98.3 % 2.2%. While the performance of the *Gene&Count* and *Gene&Binary&Count* models were nearly equivalent (98.2% ± 2.3% and 98.3% ± 2.2%, respectively), the addition of the “binary” predictor to the *Gene* model conferred some information on the resistance relationship, although it is mostly eclipsed by the more pertinent and predictive information conferred by “count”. It should be noted that on the training set, the *Gene&Binary&Count* model achieved a mean F_1_ of 98.6% ± 2.6%. The improvement in performance over single mechanism models (*Binary, Count*, and *Gene*) confirms that AMR genes do not contribute equally to antibiotic resistance. We also fit a *Full* model that considered all identified AMR genes, not exclusively those expected to confer resistance to the model antibiotic, and found the additional predictors did not lead to improved performance over the *Gene&Count* combination (F_1_ of 88.7% *±* 14.4%).

Across all evaluated antibiotics and breakpoints, the *Count* model (i.e. number of genes) demonstrated the strongest improvement in phenotype prediction compared with additional genes or the *Binary* model. The *Count* model achieved performance exceeding more complex models using only a single predictor. Surprisingly, the performance of the *Count* model suggests the number of resistance genes is more important for predicting resistance than specific gene identity, and individual genes alone cannot predict resistance. At the same time, it is also possible the *Count* model offers a low-dimensional approximation to a more complex resistance relationship involving multiple genes and their interactions that cannot be supported by the limited data in this study. Still, there were several antibiotics where the “count” variable was insufficient to predict resistance by itself. Both the *Count* and *Gene* models achieved F_1_ scores *≤* 76.9 for amoxicillin/clavulanic acid, ampicillin, and cephalexin. Notably, the joint *Gene&Count* model achieved improved F_1_ scores of ≤ 95.7 for the same antibiotics.

All models included animal main effects and animal×gene interaction effects to allow different levels and types of genetic associations across animal host. To evaluate the role of animal effects in these models, we removed animal main and interaction effects from the best performing *Gene&Binary&Count* model. The performance of the *Gene&Binary&Count* model without animal effects was nearly equivalent with an achieved mean F_1_ of 98.0% ±2.4%. Despite our initial assumptions, the inclusion of animal predictors did not impact resistance prediction on average. The lack of animal host influence on AMR resistance could be explained by the isolation of animal hosts from each other, thus limiting cross-species transmission of AMR genes. The co-linearity between different AMR genes and animal hosts would allow genetic predictors to effectively substitute for animal effects [69], but a no-animal *Count* model, which masked the identity of individual genes and did not use any animal information, achieved only a slightly lower mean F_1_ score of 90.5% *±* 14.7% compared with the animal *Count* model (mean F_1_ of 92.0%).

### Analysis of individual predictors

Because all prior results are averaged across models, it is important to understand the contribution of individual predictors to individual models and to help establish an interpretable biological understanding of AMR. We decided to explore the individual predictors in the *Gene+Binary+Count* model Although inclusion of the “binary” and animal predictors had little average effect, the elastic net handles the excess predictors with no decrease in average performance, allowing assessment of the importance of all these predictors in the individual models. We evaluated the importance, effect size, and power to detect each predictor. Given the high number of possible genetic predictors and the inevitable imbalance in predictors and response, this observational study is under-powered for many possible predictors. We therefore computed for each predictor a power to detect with a simplified model (see Methods). Across all 31 models, the average power to detect any predictor with effect size of 0.7 (two-fold increase in odds of resistance) at significance level 0.05 in the *Gene&Binary&Count* model was only 0.56 (sd = 0.22%). At an effect size of 2.3, equivalent to a ten-fold increase in the odds of resistance, the same average power was 0.90 (sd = 0.28%).

Next, we wanted estimate the relative contribution of the host animal effect. To do so we calculated the proportion of important animal-related predictors, which include the animal host predictors and their interactions with AMR genes, for each model. Of the 31 fitted models, 14 did not use any animal-related predictors, and we observed that models with fewer predictors do not consider animal information. Models with 30 or more predictors had a majority of animal-related predictors. The non-*wt* doxycycline, chloramphenicol, and neomycin were the only models with less than 27 total predictors that did not use any animal information. The models with a high proportion (≥80%) of animal interaction terms among their important predictors were non-*wt* trimethoprim-sulphamethoxazole (93.2%), cefazolin resistance (86.9%), ampicillin resistance (84.9%), non-*wt* ceftazidime (83.6%), non-*wt* amoxicillin (82.9%), gentamicin resistance (82.9%), non-*wt* spectinomycin (82%), and non-*wt* streptomycin (80%). There was a trend for non-*wt* models to select animal predictors, with 12/16 (75%) incorporating animal information, while 10/15 (67%) of resistance models did not use any. While some models used animal information, the total importance of animal predictors in the *Gene&Binary&Count* model was relatively low (Table S3), consistent with the already reported small improvement in average model performance.

### Unique predictors for resistance within an antibiotic class

Next we sought to understand the relative complexity of AMR genotype-to-phenotype relationships across different antibiotics. To do so, we compared the number of important predictors for each model, by comparing the number of important predictors was correlated with the number of candidate predictors for each antibiotic (Pearson’s correlation, *R* = 0.61). The model for amikacin and cefovecin selected a large proportion, 30/35 (85.7%) and 84/98 (85.7%), respectively, of their candidate predictors as important (Table 3). However, this pattern was not observed in other *β*-lactam class antibiotics. For ampicillin resistance, important predictors were only 26.7% of candidates, and amoxicillin-clavulanic acid resistance only selected 8/95 (8.4%) of possible predictors. The fraction of important predictors was similar for antibiotics across CLSI and ECOFF breakpoints, the model for non-*wt* ampicillin selected 12 important predictors out of 194 or 6.1%.

**Table 3.**
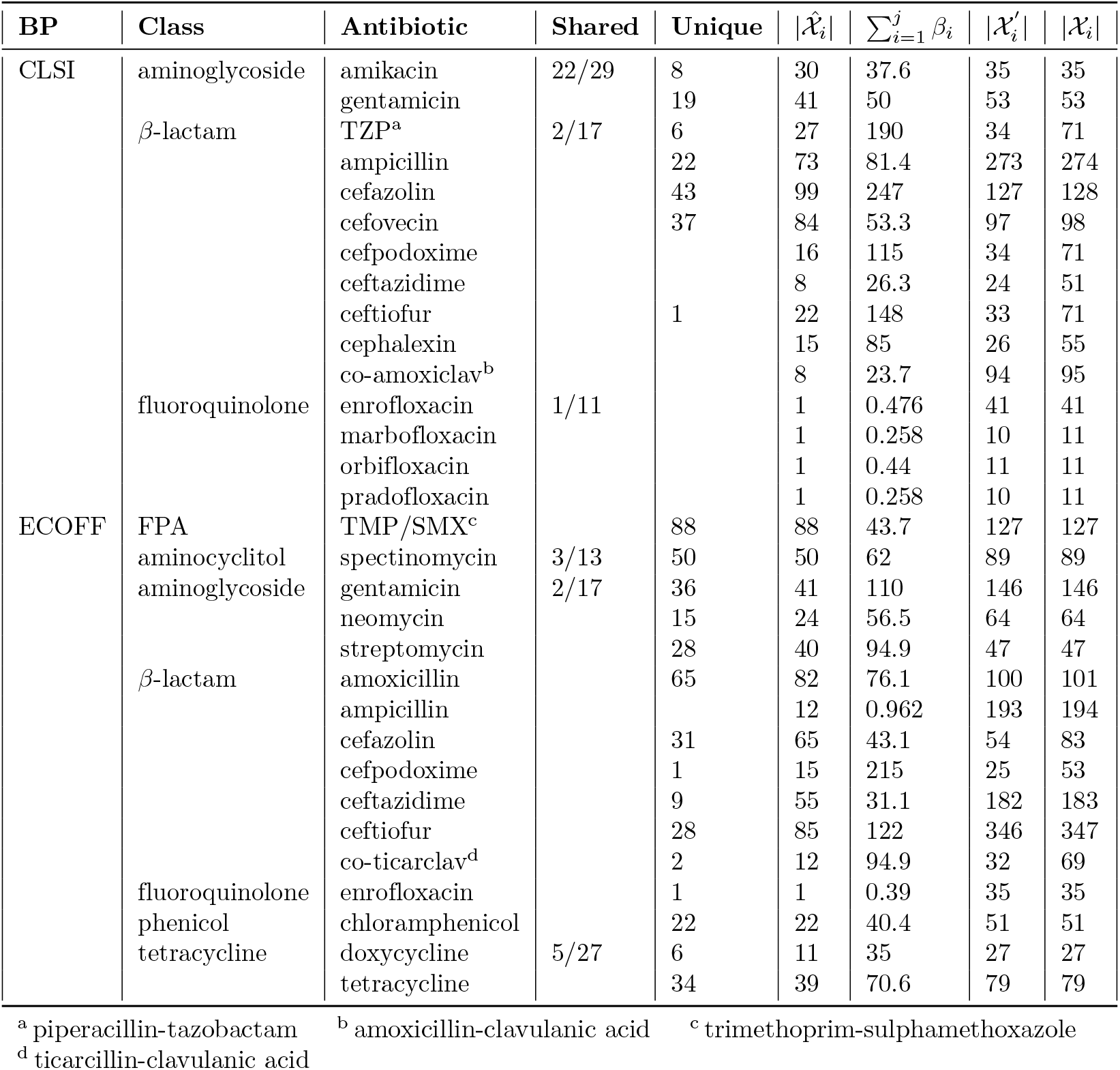
Unique predictors in fitted *Gene&Binary&Count* elastic net model by antibiotic. The important predictors included in the fitted *Gene&Binary&Count* models for each antibiotic were identified and compared to the shared important predictors for antibiotics within the same class. For each antibiotic class, “Shared” is the number of important predictors selected in all antibiotics within the same class, the denominator indicates the total number shared predictors. 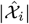 is the total number of important predictors selected by the model. 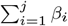 is the sum of coefficient absolute values for predictors. | χ_*i*_ | refers to the number of candidate predictors considered. 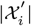 refers to the number of candidate predictors with estimated power *≥* 0.80 at an effect size of 2.3. “Unique” is the number of important predictors | χ_*i*_ | that are unique to that antibiotic relative to its class. *β*-lactams and *β*-lactam combination antibiotics had the highest number of total and unique important predictors spread across each of the antibiotics in its class. Models for fluoroquinolone resistance only agreed on a single predictor (*Count*).

In contrast with the varying complexity of *β*-lactam models, the models for fluoroquinolones were much simpler. Despite good power to detect a role for individual genes, only the “count” predictor was consistently selected for both resistant and non-*wt* phenotype prediction. For marbofloxacin, orbifloxacin, and pradofloxacin resistance models, this may be caused by the relatively low number of candidate predictors (*n* = 11). However, the models for resistant and non-*wt* enrofloxacin have more candidate predictors comparable with several *β*-lactams yet still only select for the “count” predictor. The sharing of a single predictor across different models suggests resistance or non-*wt* phenotype prediction is more similar for fluoroquinolones than other antibiotics. Aminoglycoside resistance shared a large fraction of important predictors, 22/29 (73.3%) for amikacin and 22/41 (53.6%) for gentamicin, respectively. These similarities were not present for non-*wt* aminoglycoside phenotype prediction, where the important predictors for gentamicin, neomycin, and streptomycin were mostly unique. A table of important model predictors for each antibiotic and breakpoint, not including animal interaction predictors, is provided in the supplement (Table S4).

The difference in important predictors for resistance to antibiotics within the same class may indicate that differences do exist in the genetic associations for resistance. This difference could also be explained by insufficient data in a particular antibiotic and breakpoint subset to study the antibiotic or low power when considering smaller effect sizes.

## Discussion

The availability of sequence data has been proposed as a major tool for monitoring antibiotic resistance [70–72]. Multiple studies have used WGS to predict resistance phenotypes in pathogens such as *E. coli, Klebsiella pneumoniae, Salmonella enterica* serovar Typhimurium, and *Staphylococcus aureus* [73–75]. These studies report high concordance between a pathogen’s AMR profile and predicted resistance phenotype, and consequently affirm the potential of sequencing methods to improve AMR surveillance. In recent years, the predictive accuracy the ability to extract useful information about these relationships has increased with the development of predictive ML models for AMR [13, 76]. However, we found these AMR genotype-phenotype relationships to vary substantially depending on the animal host and antibiotic, indicating a degree of AMR-predictive complexity that has not yet been discussed. These results are supported by earlier studies which show varying rates of resistance for antibiotics across animal hosts [28, 29]. While the potential utility and use of bacterial genotype data in monitoring AMR is certain, there is a need to refine current approaches to account for the biological variation in AMR mechanisms between different host animals, antibiotics, and known AMR conferring genes.

### AMR genotype-phenotype relationships are biologically complex

We observe the largest improvement in F_1_ score from the *Gene* model to *Gene&Count* models in cephalexin, ampicillin, amoxicillin-clavulanic acid, and fluoroquinolone class antibiotic resistance. This suggests the additive effect of multiple AMR genes against those antibiotics is a more effective predictor of resistance than the identity of the genes involved. From a biomechanistic perspective, orbifloxacin, a fluoroquinolone, disrupts bacterial cells by interfering with DNA replication enzymes such as topoisomerase and DNA gyrase. One mechanism of fluoroquinolone resistance is the synthesis of proteins to competitively inhibit DNA gyrase and prevent fluoroquinolones from disrupting its function [77]. For these antibiotics, the number of genes and, in turn, the number of synthesized proteins, may be a more accurate predictor of resistance than the identity of individual genes. In which case, transcriptomic data may be invaluable for fluoroquinoline resistance prediction. Another possibility is that the *Gene* model is over-parameterized in these data, and a more parsimonious model with a single predictor could achieve better performance, but in most cases the power to detect individual genes in quite high. Controlled experiments could confirm the importance of the “count” predictor, or additional samples may yet yield the statistical power to reveal the importance of gene identities. Furthermore, while the “count” of relevant AMR genes is an important predictor of resistance and models including the “count” predictor exceed the performance of the rule-based model (*Binary*) for certain antibiotics, the more complex joint model using the genomic, animal host, count, and binary predictors (*Gene&Binary&Count*) performs better still.

The genetic complexity of antibiotic resistance, in terms of the number of important predictors, was much higher for *β*-lactam and *β*-lactam combination antibiotics. The *β*-lactam class models, using both CLSI and ECOFF values, selected the highest number of unique predictors per antibiotic. More specifically, the most complex resistance models were for cefazolin, cefovecin and ampicillin. Although many of the important predictors in these models correspond to subclasses of the *β*-lactamase gene, their significance as unique predictors suggests they are non-substitutive and have varying importance in conferring resistance. Differences in *β*-lactamase protein target affinity and degree of membrane permeability may make certain subclasses of *β*-lactamase proteins more suited to act on specific antibiotics [78]. The model for resistance to amoxicillin-clavulanic acid, a different *β*-lactam class antibiotic, only selected for eight important predictors. This difference in complexity may be due to unique properties of amoxicillin-clavulanic acid or genetic mechanisms unique to amoxicillin-clavulanic acid resistance.

The difference in model complexity for each antibiotic type suggests there are antibiotic-specific AMR genotype-to-phenotype complex relationships more complex than have been previously explored [23–25]. While WGS information can be used to accurately predict AMR phenotype, the most effective way to use that information may differ between antibiotics. As an example, there were significant differences in the odds of resistance between different animal hosts for some antibiotics, and animal predictors were used to improve the predictive power of our models. Yet for other antibiotics, such as fluoroquinolones, our results show animal predictors were not necessary to make accurate predictions. In addition, the choice of specific important predictor selection in our model was significantly affected by the power to detect those predictors. Animal-specific fluoroquinolone interactions may affect resistance at a lower effect size than what is detectable in our study.

Notably, we observed a difference in the importance of selection for animal predictors in non-*wt* and resistant phenotypes. Animal predictors and interactions are more commonly found in models predicting resistance, suggesting that the animal/tissue differences respected by CLSI breakpoints also associate with differences in AMR genetic determinants. As expected, ECOFF-determined *wt* phenotypes, which are host independent, yield fewer important animal predictors. On the other hand, the development of CLSI breakpoints do take into consideration the presence/absence of numerous clinical factors, including animal host species, source tissue, and, in turn, antimicrobial- and animal-specific pharmacokinetics.

Our observations suggest that using the elastic net approach with animal predictors can accurately predict antibiotic non-*wt* phenotypes, even when those phenotypes are established using inherently non-clinical ECOFF values. As a result, the work shown here would suggest that specific antibiotics: namely, chloramphenicol, doxycycline, tetracycline, trimethoprim sulfamthoxazole, amoxicillin, neomycin, and spectinomycin, be prioritized for CLSI breakpoint analysis and development.

### AMR prediction is accurate, but incomplete

Both the *Gene&Count* and *Gene&Binary&Count* models were able to accurately predict isolate resistance. Although performance varied between different antibiotics, mean the F_1_ for both models were very high (0.988). All models share a higher average positive predictive value than negative predictive value, suggesting there are types of AMR which are not accounted for in the model and would improve negative prediction. This same complexity may also increase the chance that important resistance-conferring genes are missing from the tested predictors [79]. Theoretical power calculations for elastic net models have yet to be defined, but calculations based on traditional logistic regression suggest that despite the size of this study, our approach was underpowered considering: (1) the number of predictors and (2) an effect size equivalent to a two-fold increase in the likelihood of resistance, even when limited to only the relevant AMR genes for each antibiotic. Therefore, important predictors of AMR would best be assessed in an experiment explicitly designed to detect the effect of multiple genetic predictors. This type of experiment would require significantly more samples than were observed here to ensure sufficient statistical power, especially when measuring smaller effect sizes.

## Conclusions

Our study illustrates the genetic and environmental complexity of AMR that is often ignored in models based on ML. We confirmed that existing genetic determinants of AMR are generally accurate predictors of antibiotic resistance across animal hosts and antibiotics. However, we find evidence for possible differences in the mechanisms of AMR across host species. Using elastic net models, which offer more interpretable features than other high dimensional models, we find the genotype/phenotype relationship for AMR depends on multiple factors, including the genetic association with AMR, animal host, antibiotic, and AMR genes. While our study identifies strong genetic predictors of AMR resistance from observational samples, these predictors are merely a subset of the candidate predictor genes that we expect to confer resistance. It is uncertain if predictors not included in the model are unnecessary for predicting resistance, or whether we lack sufficient power to measure their effect on resistance. We suggest these results be further elucidated in a more powerful study, where smaller effect sizes can be detected. the effect size of individual genes can more accurately be measured. It might also be possible to test some of the biological mechanisms revealed from our interpretable elastic net models. While there is considerable evidence that WGS data are highly predictive of AMR phenotype and will improve AMR prevention strategies, we suggest there is still much to learn about the genotype-to-phenotype relationships and their differences across hosts and antibiotics. Models fit to data from multiple hosts and antibiotics may be most useful when there are commonalities in resistance mechanisms across hosts and antibiotics (e.g. in the same class), but our results show such models must also allow for some differences. In the end, such models may improve early diagnosis across hosts, and help us take action to combat resistance, especially when resistance patterns that also threaten human health emerge in non-human hosts.

## Author Declarations

### Data Availability Statement

Raw sequences used to produce the AMR output data are publicly available on NCBI’s Sequence Read Archive under BioProject ID PRJNA510384 (https://www.ncbi.nlm.nih.gov/bioproject/510384). The code and data are available at https://www.github.com/FriedbergLab/AMREcoli and on Figshare at https://doi.org/10.6084/m9.figshare.21737288.v1.

### Author Contributions

HCC, CLF, IF, KSD, and BH designed the research plan and experimental schema; HCC, CLF, JAH, TS, and BH conducted the experiments and acquired data for research analysis; all authors analyzed and interpreted the data. All authors also drafted, critically revised, and reviewed the manuscript for important intellectual content and agree to be held accountable to the accuracy and integrity of all work represented here.

## Acknowledgments

We gratefully acknowledge Dr. Christina Loiacono, Dr. Kelley Black, Dr. Kristina Lantz, Keira Stuart, Kathryn Reynolds, and Jennifer Rodriguez in the NVSL for their valuable contributions to data aggregation, collection, and processing across the NAHLN participating laboratories over the past five years. We also gratefully acknowledge the NAHLN participating laboratories that contributed WGS and AST data to this project:

Auburn University Bacteriology and Mycology Diagnostic Laboratory, Auburn, AL

California Animal Health and Food Safety Laboratory System, Davis, CA

Colorado State University Veterinary Diagnostic Laboratory, Fort Collins, CO

Kissimmee Animal Disease Diagnostic Laboratory, Kissimmee, FL

Athens Veterinary Diagnostic Laboratory, Athens, GA

Iowa State University Veterinary Diagnostic Laboratory, Ames, IA

Purdue University Animal Disease Diagnostic Laboratory, West Lafayette, IN

Kansas State University Veterinary Diagnostic Laboratory, Manhattan, KS

Murray State University, Breathitt Veterinary Center, Hopkinsville, KY

University of Kentucky Veterinary Diagnostic Laboratory, Lexington, KY

Louisiana State University Animal Disease Diagnostic Laboratory, Baton Rouge, LA

Michigan State Diagnostic Center for Population and Animal Health, Lansing, MI

University of Minnesota Veterinary Diagnostic Laboratory, MN

University of Missouri Veterinary Medical Diagnostic Laboratory, Columbia, MO

Mississippi State University Veterinary Research and Diagnostic Laboratory, Pearl, MS

North Carolina Veterinary Diagnostic Laboratory System, Rollins Animal Disease Diagnostic Laboratory, Raleigh, NC

North Dakota State University Veterinary Diagnostic Laboratory, Fargo, ND

University of Nebraska Veterinary Diagnostic Center, Lincoln, NE

Cornell Animal Health Diagnostic Center, Ithaca, NY

Ohio Department of Agriculture, Reynoldsburg, OH

Pennsylvania Animal Diagnostic Laboratory System, University of Pennsylvania New Bolton Center, Kennett Square, PA

Pennsylvania Animal Diagnostic Laboratory System, Pennsylvania Veterinary Laboratory, Harrisburg, PA

Pennsylvania State University Animal Diagnostic Laboratory, University Park, PA

South Dakota State University Animal Disease Research and Diagnostic Laboratory, Brookings, SD

Texas A&M University Veterinary Medical Diagnostic Laboratory, College Station, TX

Washington State University Animal Disease Diagnostic Laboratory, Pullman, WA

University of Wisconsin Veterinary Diagnostic Laboratory, Madison, WI

## Disclaimer

The views expressed in this article are those of the authors and do not necessarily reflect the policies and views of the USDA, DOE, or ORAU/ORISE. Mention of trade names or commercial products in this publication are solely for the purpose of providing specific information and does not imply recommendation or endorsement by the U.S. Department of Agriculture.

## Abbreviations

AAVLD: American Association of Veterinary Laboratory Diagnosticians
AMR: antibiotic resistance
APHIS: USDA Animal and Plant Health Inspection Service
AST: antibiotic sensitivity testing
BHI: brain heart infusion
CARB NAP: National Action Plan for Combating Antibiotic-Resistant Bacteria
CEAH: USDA Center for Epidemiology and Animal Health
CDC: Centers for Disease Control and Prevention co-amoxiclav, amoxicillin-clavulanic acid
co-ticarclav: ticarcillin-clavulanic acid
CLSI: Clinical Laboratory Sciences
ECOFF: epidemiological cut-off
EUCAST: European Committee on Antimicrobial Susceptibility
FDA: U.S. Food and Drug Administration
FDA-CVM-VetLIRN: FDA Center for Veterinary Medicine Veterinary Laboratory Investigation and Response Network
FPA: folate pathway antagonist
MALDI-TOF: matrix-assisted laser desorption/ionization–time of flight
MIC: minimum inhibitory concentration
ML: machine learning
NAHLN: National Animal Health Laboratory Network
NARMS: National Antibiotic Resistance Monitoring System
NCBI: National Center for Biotechnology Information
NPV: negative predictive value
NVSL: National Veterinary Services Laboratories
ORISE: Oak Ridge Institute for Science and Education
ORAU: Office of Research in Associated Universities
PPV: positive predictive value
SBA: sheep blood agar
TMP/SMX: trimethoprime-sulphamethoxazole
TrAC: Translational Artificial Intelligence Center
TSB: trypticase soy broth
TZP: piperacillin-tazobactam
USDA: U.S. Department of Agriculture
WGS: whole-genome sequencing

## Supporting information

**S1 Table. Important predictors in fitted *Gene&Binary&Count* elastic net model by antibiotic**. Listed are the important predictors identified by Gene&Binary&Count models for each antibiotic. Predictor importance is estimated by the absolute value of the predictor coefficient in the model. Animal host interaction terms are not shown.

**S2 Table. Non-interpretable phenotypes by antibiotic**. The number of sample with non-interpretable phenotypes for specific host animal and antibiotics is listed.

**S3 Table. Importance of *Gene&Binary&Count* predictors**. Predictor importance is calculated as the absolute value of the predictor coefficient in the fitted model. Importance is grouped into main effects, animal, and genetic predictors.

**S4 Table. Important predictors for *Gene&Binary&Count* model**. Listed are the important predictors identified by Gene&Binary&Count models for each antibiotic are listed. Predictor importance is estimated by the absolute value of the predictor coefficient in the model. Animal host interaction terms are not shown.

**S5 Table. *Gene&Binary&Count* model performance**. Performance metrics for the *Gene&Binary&Count* elastic net model are presented.

