## Supplemental information for "An accurate and interpretable model for antimicrobial resistance in pathogenic *Escherichia coli* from livestock and companion animal species"

**S1. Resistance phenotypes by antibiotic and breakpoint.** Listed are number of samples with each resistance phenotype separated by reference breakpoint and antibiotic. Resistance phenotypes are; I = intermediate, NI = non-interpretable, R = resistant, and S = sensitive.

| BP | Antibiotic | I | NI | R | S |
|---|---|---|---|---|---|
| CLSI | amikacin | 15 | 1 | 7 | 319 |
| | ampicillin | | 27 | 571 | 46 |
| | cefazolin | 18 | 109 | 91 | 124 |
| | cefovecin | 2 | | 26 | 105 |
| | cefpodoxime | 2 | | 54 | 138 |
| | ceftazidime | 8 | | 26 | 93 |
| | ceftiofur | | | 51 | 146 |
| | cephalexin | 29 | 18 | 97 | 49 |
| | co-amoxiclav$_a$ | 1 | 30 | 231 | 104 |
| | doxycycline* | | 93 | 52 | 3 |
| | enrofloxacin | 7 | 124 | 69 | 410 |
| | gentamicin | 5 | 1 | 48 | 288 |
| | marbofloxacin | 2 | | 41 | 256 |
| | minocycline* | | 2 | 21 | |
| | orbifloxacin | 8 | | 42 | 248 |
| | pradofloxacin | 1 | | 41 | 257 |
| | TZP$_c$ | 4 | | 9 | 180 |
| ECOFF | amikacin* | | | | 175 |
| | amoxicillin | | | 57 | 106 |
| | ampicillin | | | 91 | 82 |
| | azithromycin* | | 34 | | 116 |
| | cefazolin | | | 26 | 149 |
| | cefovecin | | 233 | | |
| | cefpodoxime | | | 19 | 153 |
| | ceftazidime | | 212 | 49 | 128 |
| | ceftiofur | | 2 | 75 | 340 |
| | cephalexin* | | 20 | | 152 |
| | cephalothin* | | | | 1 |
| | chloramphenicol | | 3 | 53 | 461 |
| | chlortetracycline | | 183 | | |
| | clarithromycin | | 151 | | |
| | clindamycin | | 464 | | |
| | danofloxacin | | 300 | | |
| | doxycycline | | | 69 | 300 |
| | enrofloxacin | | 30 | 94 | 246 |
| | erythromycin | | 315 | | |

| | | | |
|---|---|---|---|
| florfenicol* | 98 | | 365 |
| gamithromycin | 146 | | |
| gentamicin | 17 | 111 | 510 |
| imipenem* | 511 | 4 | 2 |
| marbofloxacin | 67 | | |
| minocycline* | | | 1 |
| neomycin | | 126 | 337 |
| nitrofurantoin* | | | 1 |
| novobiocin | 163 | | |
| orbifloxacin | 67 | | |
| oxacillin | 152 | | |
| oxytetracycline | 346 | | |
| penicillin | 615 | | |
| pradofloxacin | 67 | | |
| rifampin | 152 | | |
| spectinomycin | 28 | 100 | 335 |
| streptomycin | | 44 | 119 |
| sulphadimethoxine | 463 | | |
| sulphathiazole | 163 | | |
| tetracycline | 2 | 270 | 554 |
| tiamulin | 300 | | |
| ticarcillin | 146 | | |
| ticarcillin clavulanic acid | 4 | 21 | 121 |
| tildipirosin | 146 | | |
| tilmicosin | 300 | | |
| TMP/SMX$_b$ | 189 | 205 | 552 |
| tulathromycin | 271 | | |
| tylosin tartrate | 434 | | |
| TZP$_c$* | | 5 | 167 |
| vancomycin | 1 | | |

[a] amoxicillin-clavulanic acid [b] trimethoprim-sulphamethoxazole [c] piperacillin-tazobactam
* Excluded from analysis due to low sample size.

**S2. Antibiotic and Host Animals without ECOFF or CLSI breakpoints.** Listed are the number of antibiotic and host animal combinations that did not have interpretable phenotypes under CLSI or ECOFF breakpoints at the time of writing.

| Antibiotic | Host Animal | n |
|---|---|---|
| cefovecin | cat, dog | 235 |
| chlortetracycline | cattle, swine | 183 |
| clarithromycin | horse | 151 |
| clindamycin | cattle, swine, chicken, turkey, dog | 465 |
| danofloxacin | cattle, swine | 300 |
| enrofloxacin | swine | 20 |
| erythromycin | chicken, turkey, horse, dog | 316 |
| gamithromycin | cattle, swine | 146 |
| marbofloxacin | cat | 67 |
| novobiocin | chicken, turkey | 164 |
| orbifloxacin | cat | 67 |
| oxacillin | horse, dog | 152 |
| oxytetracycline | cattle, swine, chicken, turkey | 347 |
| penicillin | cattle, swine, chicken, turkey, horse, dog | 616 |
| pradofloxacin | cat | 67 |
| rifampin | horse, dog | 152 |
| sulphadimethoxine | cattle, swine, chicken, turkey | 464 |
| sulphathiazole | chicken, turkey | 164 |
| tiamulin | cattle, swine | 300 |
| ticarcillin | horse | 146 |
| tildipirosin | cattle, swine | 146 |
| tilmicosin | cattle, swine | 300 |
| tulathromycin | cattle, swine | 271 |
| tylosin tartrate | cattle, swine, chicken, turkey | 435 |
| vancomycin | dog | 1 |

**S3. Predictor importance by type.** Listed are the summed importance of Gene&Binary&Count predictors by type. "Animal All" refers to host animal main effects with any interaction terms. "Gene All" refers to main gene, binary, and count predictors.

| BP | Antibiotic | Gene Main | Animal Main | Binary | Count | Interactions | Animal All | Gene All |
|---|---|---|---|---|---|---|---|---|
| CLSI | amikacin | 16 | | 1.2 | 20 | 0.37 | 0.37 | 37 |
| | co-amoxiclav$_a$ | 11 | | 1.9 | 11 | | | 23 |
| | ampicillin | 28 | 0.076 | 3.7 | 46 | 3.7 | 3.8 | 78 |
| | cefazolin | 120 | 0.52 | 0.48 | 120 | 0.012 | 0.53 | 230 |
| | cefovecin | 27 | 0.34 | 2.9 | 20 | 0.69 | 1 | 50 |
| | cefpodoxime | 74 | | 3.2 | 35 | | | 110 |
| | ceftazidime | 9.6 | | 1.3 | 5.7 | | | 17 |
| | ceftiofur | 91 | | 3.3 | 42 | | | 140 |
| | cephalexin | 50 | | 2 | 30 | | | 82 |
| | enrofloxacin | | | | 0.48 | | | 0.48 |
| | gentamicin | 4.8 | | | 27 | 0.54 | 0.54 | 32 |
| | marbofloxacin | | | | 0.26 | | | 0.26 |
| | orbifloxacin | | | | 0.44 | | | 0.44 |
| | TZP$_c$ | 140 | | 4.7 | 27 | | | 170 |
| | pradofloxacin | | | | 0.26 | | | 0.26 |
| ECOFF | amoxicillin | 47 | | 3.3 | 24 | | | 75 |
| | ampicillin | | | | 0.96 | | | 0.96 |
| | cefazolin | 17 | 0.0054 | 2.4 | 21 | 2.2 | 2.2 | 40 |
| | cefpodoxime | 73 | | 3 | 90 | | | 170 |
| | ceftazidime | 9.8 | 0.069 | | 15 | 2.5 | 2.5 | 25 |
| | ceftiofur | 65 | | 1.2 | 34 | | | 100 |
| | chloramphenicol | 13 | 6.1 | 0.49 | 19 | 0.14 | 6.3 | 32 |
| | doxycycline | 8.9 | | 0.82 | 25 | | | 35 |
| | enrofloxacin | | | | 0.39 | | | 0.39 |
| | gentamicin | 46 | 0.092 | | 54 | 0.25 | 0.34 | 100 |
| | neomycin | 26 | 0.09 | | 30 | 1.1 | 1.2 | 55 |
| | spectinomycin | 25 | 0.13 | | 32 | 0.73 | 0.86 | 57 |
| | streptomycin | 35 | 0.14 | 0.54 | 51 | 0.033 | 0.18 | 87 |
| | tetracycline | 35 | 0.43 | 2.3 | 30 | 2.4 | 2.8 | 67 |
| | co-ticarclav | 31 | | 1.8 | 34 | | | 67 |
| | TMP/SMX$_b$ | 6.9 | 0.74 | 2.3 | 20 | 12 | 13 | 29 |

[a] amoxicillin-clavulanic acid [b] trimethoprim-sulphamethoxazole [c] piperacillin-tazobactam

**S4. Important predictors in fitted *Gene&Binary&Count* elastic net model by antibiotic.** Listed are the important predictors identified by Gene&Binary&Count models for each antibiotic are listed. Predictor importance is estimated by the absolute value of the predictor coefficient in the model. Animal host interaction terms are not shown.

| BP | Antibiotic | Important Predictors |
|---|---|---|
| CLSI | amikacin | AAC(3)-VIa, AAC(6')-Ib-cr, AAC(6')-Ib-cr5, AAC(6')-Ib', AAC(6')-Ib4, AAC(6')-IIc, binary, count |
| | co-amoxiclav$_a$ | binary, blaEC, blaTEM-1, blaTEM-1B, blaTEM-1C, blaTEM-235, count, CTX-M-14 |
| | ampicillin | binary, blaEC, blaEC-13, blaEC-18, blaEC-19, blaEC-5, blaEC-8, blaTEM-1, blaTEM-1B, CMY-2, count, dog |
| | cefazolin | binary, blaEC, blaEC-13, blaEC-18, blaEC-19, blaEC-5, blaEC-8, blaTEM-1, blaTEM-150, blaTEM-1A, blaTEM-1B, CMY-2, count, dog, horse |
| | cefovecin | binary, blaEC, blaEC-13, blaEC-18, blaEC-19, blaEC-5, blaEC-8, blaTEM-1, blaTEM-104, blaTEM-105, blaTEM-150, blaTEM-1A, blaTEM-1B, blaTEM-1C, blaTEM-235, cat, CMY-2, count, CTX-M-14, dog |
| | cefpodoxime | binary, blaEC, blaEC-13, blaEC-18, blaEC-19, blaEC-5, blaEC-8, blaTEM-1, blaTEM-102, blaTEM-104, blaTEM-150, blaTEM-1A, blaTEM-1B, CMY-2, count, CTX-M-116 |
| | ceftazidime | binary, blaEC-15, blaEC-5, blaTEM-102, blaTEM-104, CMY-2, count, CTX-M-15 |
| | ceftiofur | binary, blaEC, blaEC-13, blaEC-15, blaEC-18, blaEC-5, blaEC-8, blaTEM, blaTEM-1, blaTEM-102, blaTEM-105, blaTEM-150, blaTEM-1A, blaTEM-1C, blaTEM-235, CMY-2, count, CTX-M-15, CTX-M-55, OXA-1, OXA-395, ROB-1 |
| | cephalexin | binary, blaEC, blaEC-13, blaEC-19, blaEC-5, blaEC-8, blaTEM, blaTEM-1, blaTEM-104, blaTEM-105, blaTEM-150, blaTEM-1A, blaTEM-1B, CMY-2, count |
| | enrofloxacin | count |
| | gentamicin | AAC(3)-Iid, AAC(3)-IId, AAC(3)-VIa, AAC(6')-Ib', AAC(6')-Ib4, ANT(2")-Ia, count |
| | marbofloxacin | count |
| | orbifloxacin | count |
| | TZP$_c$ | binary, blaEC, blaEC-13, blaEC-15, blaEC-18, blaEC-19, blaEC-5, blaEC-8, blaTEM-1, blaTEM-102, blaTEM-150, blaTEM-1A, blaTEM-1B, CARB-2, CMY-107, CMY-130, CMY-2, count, CTX-M-116, CTX-M-14, CTX-M-15, CTX-M-27, CTX-M-55, OXA-1, ROB, ROB-1, ROB-2 |
| | pradofloxacin | count |
| ECOFF | amoxicillin | binary, blaEC, blaEC-13, blaEC-15, blaEC-18, blaEC-19, blaEC-5, blaEC-8, blaTEM-1, blaTEM-1B, chicken, CMY-2, count, OXA-60, OXA-60b, turkey |
| | ampicillin | binary, blaEC, blaEC-13, blaEC-15, blaEC-18, blaEC-19, blaTEM-1, blaTEM-102, blaTEM-141, CMY-2, count, CTX-M-1 |
| | cefazolin | binary, blaEC, blaEC-19, blaEC-8, blaTEM-1, blaTEM-104, blaTEM-150, blaTEM-1A, blaTEM-1B, blaTEM-34, cat, CMY-2, count, horse, OXA-364, OXA-787 |
| | cefpodoxime | binary, blaEC, blaEC-18, blaEC-19, blaEC-5, blaEC-8, blaTEM-1, blaTEM-1C, blaTEM-235, CMY-2, count, CTX-M-14, CTX-M-15, OXA-364, OXA-787 |
| | ceftazidime | blaTEM-1, blaTEM-150, blaTEM-1A, CMY-2, count, CTX-M-1, CTX-M-15, CTX-M-55, horse, SHV-12 |
| | ceftiofur | binary, blaA_Mtub, blaEC, blaEC-13, blaEC-15, blaEC-18, blaEC-19, blaEC-5, blaR1, blaTEM-1, blaTEM-102, blaTEM-12, blaTEM-141, blaTEM-150, blaTEM-1A, blaTEM-1B, blaTEM-1C, blaTEM-1D, blaTEM-235, blaZ, blaZ01, blaZ8, CARB-2, chicken, CMY-2, count, CTX-M-1, CTX-M-55, HER-3, HERA-3, horse, OXA-60, OXA-60b, PDC-10, PDC-113, SHV-12 |
| | chloramphenicol | binary, catA1, catB3, cmlA, count, floR, oqxB9 |
| | doxycycline | binary, count, tet(A), tet(B), tet34 |
| | enrofloxacin | count |
| | gentamicin | AAC(3)-Ib, AAC(3)-II, AAC(3)-Iid, AAC(3)-IId, AAC(3)-IIe, AAC(3)-IIg, AAC(3)-VIa, AAC(6')-Ib4, cat, count, swine |

| | |
|---|---|
| neomycin | AAC(3)-IV, AAC(3)-IVa, AAC(3)-VIa, APH(3')-Ia, APH(3')-IIa, APH(3')-IIb, binary, chicken, count |
| spectinomycin | aadA1, aadA10, aadA12, aadA2, aadA5, ANT(3")-Ia, ant(3")-Ih/aac(6')-IId, ANT(3")-Ii-AAC(6')-IId, cattle, count, swine, turkey |
| streptomycin | AAC(3)-VIa, aadA1, aadA2, ANT(3")-Ia, APH(3")-Ib, APH(6)-Id, binary, chicken, count, turkey |
| tetracycline | binary, cattle, chicken, count, dog, otr(A), tet(A), tet(B), tet(C), tet(M), tet34, turkey |
| co-ticarclav | binary, blaEC-13, blaEC-19, blaI, blaTEM-1, blaTEM-105, blaTEM-1B, CMY-2, count, CTX-M-1, CTX-M-15, CTX-M-55 |
| TMP/SMX$_b$ | binary, cat, cattle, chicken, count, dfrA1, dfrA12, dfrA17, dfrA5, horse, swine, turkey |

[a] amoxicillin-clavulanic acid [b] trimethoprim-sulphamethoxazole [c] piperacillin-tazobactam

**S5. Evaluation metrics for all models by antibiotic and breakpoint.** Listed are the performance metrics for all models on the test set. Values are separated by breakpoint, antibiotic, and model.

| BP | Antibiotic | Model | Specificity | Sensitivity | PPV | NPV | Accuracy | F1 |
|---|---|---|---|---|---|---|---|---|
| CLSI | amikacin | Gene | 1 | 0.6 | 1 | 0.069 | 0.61 | 0.75 |
| CLSI | co-amoxiclav | Gene | 0.38 | 0.81 | 0.37 | 0.82 | 0.51 | 0.51 |
| CLSI | ampicillin | Gene | 0.8 | 0.92 | 0.33 | 0.99 | 0.81 | 0.49 |
| CLSI | cefazolin | Gene | 0.77 | 0.96 | 0.83 | 0.94 | 0.87 | 0.89 |
| CLSI | cefovecin | Gene | 1 | 0.9 | 1 | 0.75 | 0.93 | 0.95 |
| CLSI | cefpodoxime | Gene | 0.83 | 1 | 0.93 | 1 | 0.95 | 0.97 |
| CLSI | ceftazidime | Gene | 1 | 0.95 | 1 | 0.87 | 0.96 | 0.97 |
| CLSI | cephalexin | Gene | 0.5 | 0.7 | 0.35 | 0.81 | 0.56 | 0.47 |
| CLSI | enrofloxacin | Gene | 0.88 | 0.59 | 0.96 | 0.29 | 0.63 | 0.73 |
| CLSI | gentamicin | Gene | 0.82 | 1 | 0.97 | 1 | 0.97 | 0.98 |
| CLSI | marbofloxacin | Gene | 1 | 0.31 | 1 | 0.2 | 0.41 | 0.47 |
| CLSI | orbifloxacin | Gene | 1 | 0.46 | 1 | 0.27 | 0.55 | 0.63 |
| CLSI | TZP | Gene | 1 | 0.82 | 1 | 0.13 | 0.82 | 0.9 |
| CLSI | pradofloxacin | Gene | 0.89 | 0.33 | 0.94 | 0.19 | 0.41 | 0.49 |
| CLSI | ceftiofur | Gene | 0.82 | 0.97 | 0.94 | 0.9 | 0.93 | 0.95 |
| ECOFF | ampicillin | Gene | 0.79 | 0.94 | 0.8 | 0.94 | 0.86 | 0.86 |
| ECOFF | cefazolin | Gene | 0.83 | 0.67 | 0.95 | 0.33 | 0.69 | 0.78 |
| ECOFF | cefpodoxime | Gene | 0.5 | 0.97 | 0.94 | 0.67 | 0.91 | 0.95 |
| ECOFF | ceftazidime | Gene | 0.8 | 1 | 0.93 | 1 | 0.94 | 0.96 |
| ECOFF | chloramphenicol | Gene | 0.91 | 0.96 | 0.99 | 0.71 | 0.95 | 0.97 |
| ECOFF | doxycycline | Gene | 0.86 | 0.97 | 0.97 | 0.86 | 0.95 | 0.97 |
| ECOFF | enrofloxacin | Gene | 0.42 | 0.88 | 0.8 | 0.57 | 0.75 | 0.84 |
| ECOFF | gentamicin | Gene | 0.96 | 0.96 | 0.99 | 0.85 | 0.96 | 0.98 |
| ECOFF | tetracycline | Gene | 0.94 | 0.96 | 0.97 | 0.93 | 0.96 | 0.97 |
| ECOFF | TMP/SMX | Gene | 0.93 | 1 | 0.97 | 1 | 0.98 | 0.99 |
| ECOFF | amoxicillin | Gene | 0.92 | 0.95 | 0.95 | 0.92 | 0.94 | 0.95 |
| ECOFF | ceftiofur | Gene | 0.87 | 0.97 | 0.97 | 0.87 | 0.95 | 0.97 |
| ECOFF | neomycin | Gene | 0.77 | 1 | 0.92 | 1 | 0.94 | 0.96 |
| ECOFF | spectinomycin | Gene | 0.95 | 0.94 | 0.98 | 0.83 | 0.94 | 0.96 |
| ECOFF | streptomycin | Gene | 0.89 | 0.92 | 0.96 | 0.8 | 0.91 | 0.94 |
| ECOFF | co-ticarclav | Gene | 0.4 | 0.88 | 0.88 | 0.4 | 0.8 | 0.88 |
| CLSI | amikacin | Gene&Binary | 1 | 0.6 | 1 | 0.069 | 0.61 | 0.75 |
| CLSI | co-amoxiclav | Gene&Binary | 0.64 | 0.81 | 0.5 | 0.88 | 0.69 | 0.62 |
| CLSI | ampicillin | Gene&Binary | 0.91 | 0.75 | 0.47 | 0.97 | 0.9 | 0.58 |
| CLSI | cefazolin | Gene&Binary | 0.82 | 0.96 | 0.86 | 0.95 | 0.89 | 0.91 |
| CLSI | cefovecin | Gene&Binary | 0.83 | 0.95 | 0.95 | 0.83 | 0.93 | 0.95 |
| CLSI | cefpodoxime | Gene&Binary | 0.67 | 1 | 0.88 | 1 | 0.9 | 0.93 |
| CLSI | ceftazidime | Gene&Binary | 1 | 0.95 | 1 | 0.87 | 0.96 | 0.97 |
| CLSI | cephalexin | Gene&Binary | 0.73 | 0.8 | 0.53 | 0.9 | 0.75 | 0.64 |
| CLSI | enrofloxacin | Gene&Binary | 0.88 | 0.59 | 0.96 | 0.29 | 0.63 | 0.73 |
| CLSI | gentamicin | Gene&Binary | 0.82 | 1 | 0.97 | 1 | 0.97 | 0.98 |
| CLSI | marbofloxacin | Gene&Binary | 1 | 0.31 | 1 | 0.2 | 0.41 | 0.47 |
| CLSI | orbifloxacin | Gene&Binary | 1 | 0.46 | 1 | 0.27 | 0.55 | 0.63 |
| CLSI | TZP | Gene&Binary | 1 | 0.82 | 1 | 0.13 | 0.82 | 0.9 |
| CLSI | pradofloxacin | Gene&Binary | 0.89 | 0.33 | 0.94 | 0.19 | 0.41 | 0.49 |
| CLSI | ceftiofur | Gene&Binary | 0.82 | 0.97 | 0.94 | 0.9 | 0.93 | 0.95 |
| ECOFF | ampicillin | Gene&Binary | 0.79 | 1 | 0.81 | 1 | 0.89 | 0.89 |
| ECOFF | cefazolin | Gene&Binary | 0.83 | 0.6 | 0.95 | 0.29 | 0.64 | 0.73 |
| ECOFF | cefpodoxime | Gene&Binary | 0.5 | 1 | 0.94 | 1 | 0.94 | 0.97 |
| ECOFF | ceftazidime | Gene&Binary | 0.8 | 1 | 0.93 | 1 | 0.94 | 0.96 |
| ECOFF | chloramphenicol | Gene&Binary | 0.91 | 1 | 0.99 | 1 | 0.99 | 0.99 |
| ECOFF | doxycycline | Gene&Binary | 0.86 | 0.98 | 0.97 | 0.92 | 0.96 | 0.98 |
| ECOFF | enrofloxacin | Gene&Binary | 0.16 | 1 | 0.76 | 1 | 0.77 | 0.86 |
| ECOFF | gentamicin | Gene&Binary | 0.96 | 0.99 | 0.99 | 0.96 | 0.98 | 0.99 |
| ECOFF | tetracycline | Gene&Binary | 0.98 | 0.98 | 0.99 | 0.96 | 0.98 | 0.99 |
| ECOFF | TMP/SMX | Gene&Binary | 0.95 | 1 | 0.98 | 1 | 0.99 | 0.99 |
| ECOFF | amoxicillin | Gene&Binary | 0.92 | 0.95 | 0.95 | 0.92 | 0.94 | 0.95 |
| ECOFF | ceftiofur | Gene&Binary | 0.87 | 0.97 | 0.97 | 0.87 | 0.95 | 0.97 |
| ECOFF | neomycin | Gene&Binary | 0.81 | 1 | 0.93 | 1 | 0.95 | 0.96 |
| ECOFF | spectinomycin | Gene&Binary | 1 | 0.87 | 1 | 0.69 | 0.9 | 0.93 |
| ECOFF | streptomycin | Gene&Binary | 0.89 | 0.96 | 0.96 | 0.89 | 0.94 | 0.96 |
| ECOFF | co-ticarclav | Gene&Binary | 0.4 | 0.88 | 0.88 | 0.4 | 0.8 | 0.88 |
| CLSI | amikacin | Gene&Count | 1 | 0.97 | 1 | 0.5 | 0.97 | 0.98 |
| CLSI | co-amoxiclav | Gene&Count | 1 | 1 | 1 | 1 | 1 | 1 |
| CLSI | ampicillin | Gene&Count | 1 | 0.92 | 1 | 0.99 | 0.99 | 0.96 |
| CLSI | cefazolin | Gene&Count | 0.95 | 1 | 0.96 | 1 | 0.98 | 0.98 |
| CLSI | cefovecin | Gene&Count | 1 | 0.95 | 1 | 0.86 | 0.96 | 0.98 |
| CLSI | cefpodoxime | Gene&Count | 1 | 0.93 | 1 | 0.86 | 0.95 | 0.96 |
| CLSI | ceftazidime | Gene&Count | 1 | 0.95 | 1 | 0.87 | 0.96 | 0.97 |
| CLSI | cephalexin | Gene&Count | 1 | 1 | 1 | 1 | 1 | 1 |
| CLSI | enrofloxacin | Gene&Count | 1 | 1 | 1 | 1 | 1 | 1 |
| CLSI | gentamicin | Gene&Count | 0.91 | 1 | 0.98 | 1 | 0.99 | 0.99 |
| CLSI | marbofloxacin | Gene&Count | 1 | 1 | 1 | 1 | 1 | 1 |
| CLSI | orbifloxacin | Gene&Count | 1 | 1 | 1 | 1 | 1 | 1 |
| CLSI | TZP | Gene&Count | 1 | 0.97 | 1 | 0.5 | 0.97 | 0.99 |
| CLSI | pradofloxacin | Gene&Count | 1 | 0.98 | 1 | 0.9 | 0.98 | 0.99 |
| CLSI | ceftiofur | Gene&Count | 1 | 0.97 | 1 | 0.92 | 0.98 | 0.98 |
| ECOFF | ampicillin | Gene&Count | 0.79 | 1 | 0.81 | 1 | 0.89 | 0.89 |
| ECOFF | cefazolin | Gene&Count | 1 | 0.9 | 1 | 0.67 | 0.92 | 0.95 |
| ECOFF | cefpodoxime | Gene&Count | 1 | 0.97 | 1 | 0.8 | 0.97 | 0.98 |
| ECOFF | ceftazidime | Gene&Count | 0.9 | 1 | 0.96 | 1 | 0.97 | 0.98 |
| ECOFF | chloramphenicol | Gene&Count | 1 | 0.96 | 1 | 0.73 | 0.96 | 0.98 |
| ECOFF | doxycycline | Gene&Count | 1 | 0.98 | 1 | 0.93 | 0.99 | 0.99 |
| ECOFF | enrofloxacin | Gene&Count | 1 | 0.98 | 1 | 0.95 | 0.99 | 0.99 |
| ECOFF | gentamicin | Gene&Count | 1 | 1 | 1 | 1 | 1 | 1 |
| ECOFF | tetracycline | Gene&Count | 1 | 1 | 1 | 1 | 1 | 1 |
| ECOFF | TMP/SMX | Gene&Count | 1 | 1 | 1 | 1 | 1 | 1 |
| ECOFF | amoxicillin | Gene&Count | 1 | 0.95 | 1 | 0.92 | 0.97 | 0.98 |
| ECOFF | ceftiofur | Gene&Count | 1 | 0.99 | 1 | 0.94 | 0.99 | 0.99 |
| ECOFF | neomycin | Gene&Count | 1 | 1 | 1 | 1 | 1 | 1 |

| | | | | | | | | |
|---|---|---|---|---|---|---|---|---|
| ECOFF | spectinomycin | Gene&Count | 1 | 0.99 | 1 | 0.95 | 0.99 | 0.99 |
| ECOFF | streptomycin | Gene&Count | 1 | 1 | 1 | 1 | 1 | 1 |
| ECOFF | co-ticarclav | Gene&Count | 0.6 | 0.96 | 0.92 | 0.75 | 0.9 | 0.94 |
| CLSI | amikacin | Gene&Binary&Count | 1 | 0.97 | 1 | 0.5 | 0.97 | 0.98 |
| CLSI | co-amoxiclav | Gene&Binary&Count | 1 | 1 | 1 | 1 | 1 | 1 |
| CLSI | ampicillin | Gene&Binary&Count | 1 | 0.92 | 1 | 0.99 | 0.99 | 0.96 |
| CLSI | cefazolin | Gene&Binary&Count | 0.95 | 1 | 0.96 | 1 | 0.98 | 0.98 |
| CLSI | cefovecin | Gene&Binary&Count | 1 | 1 | 1 | 1 | 1 | 1 |
| CLSI | cefpodoxime | Gene&Binary&Count | 1 | 0.93 | 1 | 0.86 | 0.95 | 0.96 |
| CLSI | ceftazidime | Gene&Binary&Count | 1 | 0.95 | 1 | 0.87 | 0.96 | 0.97 |
| CLSI | cephalexin | Gene&Binary&Count | 1 | 1 | 1 | 1 | 1 | 1 |
| CLSI | enrofloxacin | Gene&Binary&Count | 1 | 1 | 1 | 1 | 1 | 1 |
| CLSI | gentamicin | Gene&Binary&Count | 0.91 | 1 | 0.98 | 1 | 0.99 | 0.99 |
| CLSI | marbofloxacin | Gene&Binary&Count | 1 | 1 | 1 | 1 | 1 | 1 |
| CLSI | orbifloxacin | Gene&Binary&Count | 1 | 1 | 1 | 1 | 1 | 1 |
| CLSI | TZP | Gene&Binary&Count | 1 | 0.95 | 1 | 0.33 | 0.95 | 0.97 |
| CLSI | pradofloxacin | Gene&Binary&Count | 1 | 0.98 | 1 | 0.9 | 0.98 | 0.99 |
| CLSI | ceftiofur | Gene&Binary&Count | 1 | 0.97 | 1 | 0.92 | 0.98 | 0.98 |
| ECOFF | ampicillin | Gene&Binary&Count | 0.79 | 1 | 0.81 | 1 | 0.89 | 0.89 |
| ECOFF | cefazolin | Gene&Binary&Count | 1 | 0.9 | 1 | 0.67 | 0.92 | 0.95 |
| ECOFF | cefpodoxime | Gene&Binary&Count | 1 | 0.97 | 1 | 0.8 | 0.97 | 0.98 |
| ECOFF | ceftazidime | Gene&Binary&Count | 0.9 | 1 | 0.96 | 1 | 0.97 | 0.98 |
| ECOFF | chloramphenicol | Gene&Binary&Count | 1 | 0.96 | 1 | 0.73 | 0.96 | 0.98 |
| ECOFF | doxycycline | Gene&Binary&Count | 1 | 0.98 | 1 | 0.93 | 0.99 | 0.99 |
| ECOFF | enrofloxacin | Gene&Binary&Count | 1 | 0.98 | 1 | 0.95 | 0.99 | 0.99 |
| ECOFF | gentamicin | Gene&Binary&Count | 1 | 1 | 1 | 1 | 1 | 1 |
| ECOFF | tetracycline | Gene&Binary&Count | 1 | 1 | 1 | 1 | 1 | 1 |
| ECOFF | TMP/SMX | Gene&Binary&Count | 1 | 1 | 1 | 1 | 1 | 1 |
| ECOFF | amoxicillin | Gene&Binary&Count | 1 | 0.95 | 1 | 0.92 | 0.97 | 0.98 |
| ECOFF | ceftiofur | Gene&Binary&Count | 1 | 0.99 | 1 | 0.94 | 0.99 | 0.99 |
| ECOFF | neomycin | Gene&Binary&Count | 1 | 1 | 1 | 1 | 1 | 1 |
| ECOFF | spectinomycin | Gene&Binary&Count | 1 | 0.97 | 1 | 0.91 | 0.98 | 0.98 |
| ECOFF | streptomycin | Gene&Binary&Count | 1 | 1 | 1 | 1 | 1 | 1 |
| ECOFF | co-ticarclav | Gene&Binary&Count | 0.8 | 0.96 | 0.96 | 0.8 | 0.93 | 0.96 |
| CLSI | amikacin | Full | 1 | 0.78 | 1 | 0.12 | 0.78 | 0.87 |
| CLSI | co-amoxiclav | Full | 0.28 | 0.81 | 0.33 | 0.76 | 0.44 | 0.47 |
| CLSI | ampicillin | Full | 0.79 | 0.92 | 0.32 | 0.99 | 0.81 | 0.48 |
| CLSI | cefazolin | Full | 0.77 | 0.88 | 0.81 | 0.85 | 0.83 | 0.85 |
| CLSI | cefovecin | Full | 1 | 1 | 1 | 1 | 1 | 1 |
| CLSI | cefpodoxime | Full | 0.83 | 1 | 0.93 | 1 | 0.95 | 0.97 |
| CLSI | ceftazidime | Full | 1 | 0.95 | 1 | 0.87 | 0.96 | 0.97 |
| CLSI | cephalexin | Full | 0.5 | 0.7 | 0.35 | 0.81 | 0.56 | 0.47 |
| CLSI | enrofloxacin | Full | 0.75 | 0.9 | 0.95 | 0.6 | 0.88 | 0.92 |
| CLSI | gentamicin | Full | 0.91 | 0.97 | 0.98 | 0.83 | 0.96 | 0.97 |
| CLSI | marbofloxacin | Full | 0.78 | 0.94 | 0.96 | 0.7 | 0.92 | 0.95 |
| CLSI | orbifloxacin | Full | 0.9 | 0.92 | 0.98 | 0.69 | 0.92 | 0.95 |
| CLSI | TZP | Full | 1 | 0.82 | 1 | 0.13 | 0.82 | 0.9 |
| CLSI | pradofloxacin | Full | 0.89 | 0.92 | 0.98 | 0.67 | 0.92 | 0.95 |
| CLSI | ceftiofur | Full | 0.82 | 0.87 | 0.93 | 0.69 | 0.85 | 0.9 |
| ECOFF | ampicillin | Full | 0.74 | 1 | 0.77 | 1 | 0.86 | 0.87 |
| ECOFF | cefazolin | Full | 0.67 | 0.77 | 0.92 | 0.36 | 0.75 | 0.84 |
| ECOFF | cefpodoxime | Full | 0.75 | 0.9 | 0.97 | 0.5 | 0.89 | 0.93 |
| ECOFF | ceftazidime | Full | 0.9 | 0.96 | 0.96 | 0.9 | 0.94 | 0.96 |
| ECOFF | chloramphenicol | Full | 0.91 | 0.91 | 0.99 | 0.56 | 0.91 | 0.95 |
| ECOFF | doxycycline | Full | 0.86 | 0.95 | 0.97 | 0.8 | 0.93 | 0.96 |
| ECOFF | enrofloxacin | Full | 0.53 | 0.96 | 0.84 | 0.83 | 0.84 | 0.9 |
| ECOFF | gentamicin | Full | 1 | 0.95 | 1 | 0.82 | 0.96 | 0.97 |
| ECOFF | tetracycline | Full | 0.94 | 0.96 | 0.97 | 0.93 | 0.96 | 0.97 |
| ECOFF | TMP/SMX | Full | 0.9 | 0.99 | 0.96 | 0.97 | 0.97 | 0.98 |
| ECOFF | amoxicillin | Full | 0.92 | 0.95 | 0.95 | 0.92 | 0.94 | 0.95 |
| ECOFF | ceftiofur | Full | 0.67 | 0.88 | 0.92 | 0.56 | 0.84 | 0.9 |
| ECOFF | neomycin | Full | 0.73 | 0.94 | 0.9 | 0.83 | 0.88 | 0.92 |
| ECOFF | spectinomycin | Full | 1 | 0.87 | 1 | 0.69 | 0.9 | 0.93 |
| ECOFF | streptomycin | Full | 0.89 | 0.92 | 0.96 | 0.8 | 0.91 | 0.94 |
| ECOFF | co-ticarclav | Full | 0.6 | 0.92 | 0.92 | 0.6 | 0.87 | 0.92 |
| CLSI | amikacin | Count | 1 | 0.93 | 1 | 0.29 | 0.93 | 0.96 |
| CLSI | co-amoxiclav | Count | 0.72 | 0.71 | 0.54 | 0.85 | 0.72 | 0.61 |
| CLSI | ampicillin | Count | 0.87 | 0.92 | 0.42 | 0.99 | 0.87 | 0.58 |
| CLSI | cefazolin | Count | 0.73 | 0.88 | 0.79 | 0.84 | 0.81 | 0.83 |
| CLSI | cefovecin | Count | 1 | 0.86 | 1 | 0.67 | 0.89 | 0.92 |
| CLSI | cefpodoxime | Count | 0.92 | 0.89 | 0.96 | 0.79 | 0.9 | 0.93 |
| CLSI | ceftazidime | Count | 1 | 0.89 | 1 | 0.78 | 0.92 | 0.94 |
| CLSI | cephalexin | Count | 0.77 | 1 | 0.63 | 1 | 0.83 | 0.77 |
| CLSI | enrofloxacin | Count | 1 | 1 | 1 | 1 | 1 | 1 |
| CLSI | gentamicin | Count | 0.91 | 1 | 0.98 | 1 | 0.99 | 0.99 |
| CLSI | marbofloxacin | Count | 1 | 1 | 1 | 1 | 1 | 1 |
| CLSI | orbifloxacin | Count | 1 | 1 | 1 | 1 | 1 | 1 |
| CLSI | TZP | Count | 1 | 0.87 | 1 | 0.17 | 0.87 | 0.93 |
| CLSI | pradofloxacin | Count | 1 | 0.98 | 1 | 0.9 | 0.98 | 0.99 |
| CLSI | ceftiofur | Count | 0.91 | 0.77 | 0.96 | 0.59 | 0.8 | 0.85 |
| ECOFF | ampicillin | Count | 0.79 | 1 | 0.81 | 1 | 0.89 | 0.89 |
| ECOFF | cefazolin | Count | 0.67 | 0.8 | 0.92 | 0.4 | 0.78 | 0.86 |
| ECOFF | cefpodoxime | Count | 0.75 | 0.94 | 0.97 | 0.6 | 0.91 | 0.95 |
| ECOFF | ceftazidime | Count | 0.9 | 0.88 | 0.96 | 0.75 | 0.89 | 0.92 |
| ECOFF | chloramphenicol | Count | 1 | 0.94 | 1 | 0.65 | 0.94 | 0.97 |
| ECOFF | doxycycline | Count | 0.86 | 0.98 | 0.97 | 0.92 | 0.96 | 0.98 |
| ECOFF | enrofloxacin | Count | 1 | 0.98 | 1 | 0.95 | 0.99 | 0.99 |
| ECOFF | gentamicin | Count | 1 | 0.96 | 1 | 0.85 | 0.97 | 0.98 |
| ECOFF | tetracycline | Count | 0.98 | 0.98 | 0.99 | 0.96 | 0.98 | 0.99 |
| ECOFF | TMP/SMX | Count | 1 | 1 | 1 | 1 | 1 | 1 |
| ECOFF | amoxicillin | Count | 1 | 0.95 | 1 | 0.92 | 0.97 | 0.98 |
| ECOFF | ceftiofur | Count | 0.8 | 0.85 | 0.95 | 0.55 | 0.84 | 0.9 |
| ECOFF | neomycin | Count | 1 | 1 | 1 | 1 | 1 | 1 |
| ECOFF | spectinomycin | Count | 1 | 0.87 | 1 | 0.69 | 0.9 | 0.93 |
| ECOFF | streptomycin | Count | 0.89 | 0.96 | 0.96 | 0.89 | 0.94 | 0.96 |
| ECOFF | co-ticarclav | Count | 0.6 | 0.92 | 0.92 | 0.6 | 0.87 | 0.92 |

| | | | | | | | | |
|---|---|---|---|---|---|---|---|---|
| CLSI | amikacin | Binary | 1 | 0.6 | 1 | 0.069 | 0.61 | 0.75 |
| CLSI | co-amoxiclav | Binary | 1 | 0.14 | 1 | 0.72 | 0.74 | 0.25 |
| CLSI | ampicillin | Binary | 0.79 | 0.92 | 0.31 | 0.99 | 0.8 | 0.47 |
| CLSI | cefazolin | Binary | 0.23 | 0.96 | 0.59 | 0.83 | 0.62 | 0.73 |
| CLSI | cefovecin | Binary | 0.5 | 0.52 | 0.79 | 0.23 | 0.52 | 0.63 |
| CLSI | cefpodoxime | Binary | 1 | 0.29 | 1 | 0.38 | 0.5 | 0.44 |
| CLSI | ceftazidime | Binary | 1 | 0.37 | 1 | 0.37 | 0.54 | 0.54 |
| CLSI | cephalexin | Binary | 1 | 0.4 | 1 | 0.81 | 0.83 | 0.57 |
| CLSI | enrofloxacin | Binary | 0.88 | 0.59 | 0.96 | 0.29 | 0.63 | 0.73 |
| CLSI | gentamicin | Binary | 0.82 | 1 | 0.97 | 1 | 0.97 | 0.98 |
| CLSI | marbofloxacin | Binary | 1 | 0.31 | 1 | 0.2 | 0.41 | 0.47 |
| CLSI | orbifloxacin | Binary | 1 | 0.46 | 1 | 0.27 | 0.55 | 0.63 |
| CLSI | TZP | Binary | 1 | 0.18 | 1 | 0.031 | 0.21 | 0.31 |
| CLSI | pradofloxacin | Binary | 0.89 | 0.33 | 0.94 | 0.19 | 0.41 | 0.49 |
| CLSI | ceftiofur | Binary | 1 | 0.13 | 1 | 0.3 | 0.37 | 0.24 |
| ECOFF | ampicillin | Binary | 0.58 | 0.71 | 0.6 | 0.69 | 0.64 | 0.65 |
| ECOFF | cefazolin | Binary | 1 | 0.27 | 1 | 0.21 | 0.39 | 0.42 |
| ECOFF | cefpodoxime | Binary | 1 | 0.48 | 1 | 0.2 | 0.54 | 0.65 |
| ECOFF | ceftazidime | Binary | 0.6 | 1 | 0.87 | 1 | 0.89 | 0.93 |
| ECOFF | chloramphenicol | Binary | 0.91 | 1 | 0.99 | 1 | 0.99 | 0.99 |
| ECOFF | doxycycline | Binary | 0.86 | 0.98 | 0.97 | 0.92 | 0.96 | 0.98 |
| ECOFF | enrofloxacin | Binary | 0.16 | 1 | 0.76 | 1 | 0.77 | 0.86 |
| ECOFF | gentamicin | Binary | 0.96 | 0.99 | 0.99 | 0.96 | 0.98 | 0.99 |
| ECOFF | tetracycline | Binary | 0.98 | 0.98 | 0.99 | 0.96 | 0.98 | 0.99 |
| ECOFF | TMP/SMX | Binary | 0.95 | 1 | 0.98 | 1 | 0.99 | 0.99 |
| ECOFF | amoxicillin | Binary | 1 | 0.5 | 1 | 0.52 | 0.68 | 0.67 |
| ECOFF | ceftiofur | Binary | 0.8 | 0.66 | 0.94 | 0.34 | 0.69 | 0.78 |
| ECOFF | neomycin | Binary | 0.81 | 1 | 0.93 | 1 | 0.95 | 0.96 |
| ECOFF | spectinomycin | Binary | 1 | 0.87 | 1 | 0.69 | 0.9 | 0.93 |
| ECOFF | streptomycin | Binary | 0.89 | 0.96 | 0.96 | 0.89 | 0.94 | 0.96 |
| ECOFF | co-ticarclav | Binary | 1 | 0.32 | 1 | 0.23 | 0.43 | 0.48 |
